# To deliberate, remember; to anticipate, forget: Cognitive deliberation profiles underpinning active forgetting-dependent everyday-like memory performance in young and aged mice

**DOI:** 10.1101/2023.04.29.538679

**Authors:** Christopher Stevens, Shaam Al Abed, Azza Sellami, Eva Ducourneau, Cathy Lacroix, Mathilde Bouchet, Faustine Roudier, Giovanni Marsicano, Aline Marighetto

## Abstract

Recalling a specific past episode that will enable us to decide which action is suited to a given present situation is a core element of everyday life. A wealth of research has demonstrated that such selective remembering is dependent upon a capacity to inhibit or provisionally ‘forget’ related yet inappropriate memory episodes which could orient behavior in unwilled directions. Everyday-like memory (EdM) refers to this type of common organizational mnemonic capacity, known to deteriorate significantly with age, putatively as a result of decline in the cognitive capacity for selective inhibition or ‘active forgetting’. Moreover, this memory retrieval-concomitant active forgetting comes at the cost of genuine amnesic weakening of the inhibited episodes, a phenomenon referred to as retrieval-induced forgetting (RIF). In the present study, we introduce a novel characterization of our previously validated mouse model of EdM in terms of the existing active forgetting and RIF literature. We also introduce novel behavioral analyses of the deliberation processes elicited by EdM challenge and use detailed multi-factorial explorations to reveal how these processes are impacted by age, temporal retention demand, difficulty of EdM challenge, and anticipation of trial outcome. Our observations indicate that deliberation requires remembering while accurate anticipation—in which a critical age-related deficit is also observed—requires active forgetting. Our results represent a significant advance towards unifying our understanding of the neurocognitive processes underpinning everyday-like memory, RIF, mnemonic deliberation, anticipatory function, and how they all are impacted by the physiological ageing process. In parallel, we present preliminary results using a transgenic mouse model which point to a fundamental role for the endocannabinoid system (eCS) in active forgetting and EdM, thereby demonstrating that deeper investigation of previously characterized age-related decline of the eCS should be a pre-clinical priority with a view to developing treatments for age-related decline of EdM function.

## 1. Introduction

The cognitive fabric of our daily lives is woven together from a range of sensory and mnemonic threads, each varying in texture and quality. The voluntary actions we choose to perform in a given spatiotemporal sensory context are informed by our organisms’ retention and selective recall of what has previously been done and experienced in the same or similar environments. Whereas classical rodent memory protocols have focused primarily on retention or working memory (WM) capacities in low-to-no interference contexts (as noted in Dumas et al., 2010), the central distinguishing feature of the hippocampus-dependent everyday-like episodic memory (EdM) model we employ here is its focus on complex organizational and selective attentional cognitive challenges reflective of real world situations (Al Abed et al., 2016; adapted from the WM task originally described in Mingaud et al., 2008; Marighetto et al., 2008). This proximity to everyday human memory demands makes the model a powerful preclinical tool, of which we have already validated a virtual version for human studies (Marighetto et al., 2011; Etchamendy et al., in prep.).

### 1.1 Everyday-like memory

The EdM model (figure 1) consists of a global environment (radial maze) within which animals are mnemonically challenged according to a pseudorandom sequence in three related but spatially distinct local task contexts (radial maze arm pairs *A, B, C*). On each trial *n* in a given local context (e.g. pair *A*), the animal must recall which arm it visited during the previous *n-1* trial on the same pair in order to choose the opposite arm and get a reward. Hence, on any given trial, only one spatially and temporally determined memory episode (i.e. ‘arm explored in this spatial context on most recent trial’) can reliably guide correct choice response (i.e. ‘explore arm not visited on most recent trial’), in accordance with the spontaneous rodent behavior of spatial alternation (Richman et al., 1986; Dember & Richman, 1989). Through variation in the number of interposed trials an animal must complete between a trial *n-1* and trial *n* on a given pair, the model gives rise to EdM challenges of varying difficulty. Successful performance therefore relies on accurate selection of spatiotemporal context-appropriate memory episodes from among an overlapping set of global and local environment-related cognitive content.

**Figure 1.**
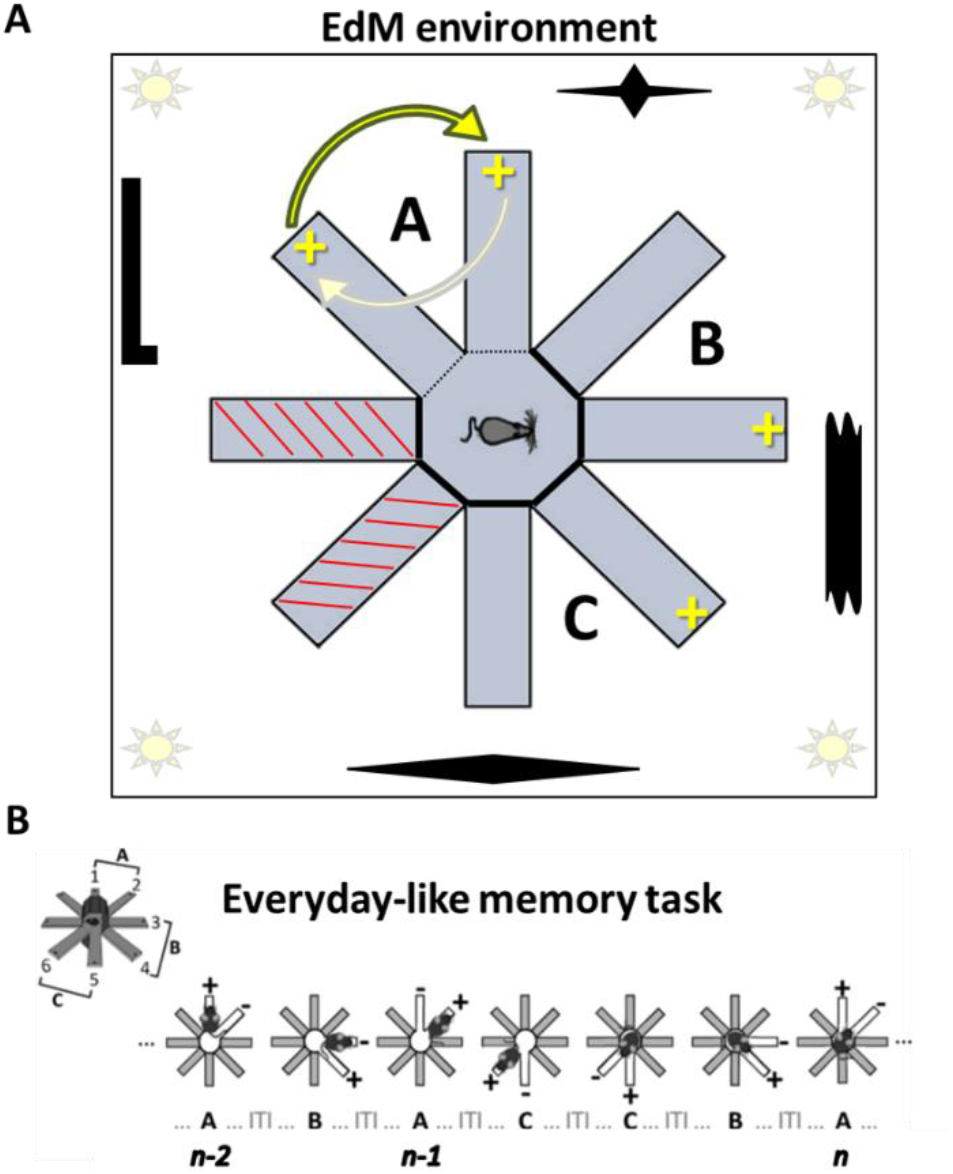
Schematic of everyday-like memory (EdM) model radial maze environment plus sample trial sequence. **A)** Six arms of the eight-arm radial maze are used, grouped into three pairs, *A, B*, and *C*. ‘Landmarks’ are placed on the walls of the radial maze room, by which animals can spatially locate themselves. Food reward distributors are located in the most distal zone of the arms. Once reward has been recuperated from one arm of a pair, it will be available in the opposite arm on the subsequent trial on that pair. **B)** Sample trial sequence from EdM protocol. Looking at trial *n* here (furthest right) we can see clearly how recall of the memory episode corresponding to *A*_*n-2*_ could lead the animal into error by making it “alternate” relative to that memory, instead of relative to the memory of *A*_*n-1*_, into the unrewarded arm. We can also see that trial *A*_*n-1*_ was a difficulty level 1 trial (one interposed trial on pairs *B* or *C* between *A*_*n-2*_ and *A*_*n-1*_) and that trial *A*_*n*_ is a difficulty level 3 trial (three interposed trials on pairs *B* or *C* between *A*_*n-1*_ and *A*_*n*_).

### 1.2 Active forgetting

Recent studies in rodents and humans have demonstrated that such memory selection processes do not consist simply in activation of the appropriate memory item but also require the concomitant *inhibition* of inappropriate but related mnemonic content which may be evoked by the global or local environment and thereby intrude into conscious attention, competing with and perturbing accurate local action choice (M. C. Anderson & Hulbert, 2021; Brewin & Smart, 2005; Wimber et al., 2015). The literature refers to such inhibition of interference as adaptive, directed, or active forgetting (M. C. Anderson & Hulbert, 2021; Tanaka et al., 2019; Costanzi et al., 2021), where this term is to be understood in the organizational and attentional sense of ‘Forget about that for a moment’ rather than as total erasure of a memory episode.

This organizational conception of memory function fits with classical theories and computational models conceiving of cognitive consciousness as a global workspace (GW) of limited capacity (Baars, 1988, 1997, 2002; Dehaene et al., 2003; Dehaene & Changeux, 2011; for a novel distributed neuronal version of the global workspace, or GNW, see Mashour et al., 2020). This workspace—where the WM functions of perceptual attention and mnemonic retrieval are theorized to play out (Lustig & Jantz, 2015)—can be pictured as a conscious cognitive meeting point where sensory input from the external environment and elicited mnemonic content from the internal (cortico-hippocampal) environment interact (for a theorization and computational model of memory as a dimension of environment, see Zilli & Hasselmo, 2008). From here on, we will use the label WM, in its deepest sense, to refer to this conscious cognitive workspace. In such a model, the function of active forgetting can be understood analogously to that of perceptual attentional processes which, by managing inhibition of potentially distracting content, grant access to WM only to a limited range of sensory input. The aim of the EdM model is to challenge organizational memory function conceived of in just this inhibition-based manner.

### 1.3 Retrieval-induced forgetting

Behind its organizational function, it has nevertheless been demonstrated that active forgetting during retrieval does also exert a lasting and genuinely amnesic effect upon mnemonic items thereby inhibited, a phenomenon referred to as retrieval-induced forgetting, or RIF (studies in humans: M. C. Anderson et al., 1994; Hulbert et al., 2016; studies in rats: Bekinschtein et al., 2018; Wu et al., 2014; Yamada et al., 2014). Hence, cognitive content subjected to RIF becomes more difficult to retrieve at a future point, if and when needed. Similar amnesic effects have previously been observed in humans following the inhibition of distracting sensory input (Biss et al., 2013). In short, forgetting or ignoring something ‘for a moment’, in order to adaptively focus on an immediate task, nevertheless seems to entail increased risk of unreliable recall of that item next time it *is* required.

### 1.4 Translational validity

We can illustrate the complex daily life implications of the above processes by representing a situation where we have to go to another part of our home or workplace in order to retrieve an item necessary for the completion of a task at hand. Upon arriving into the new environment (e.g. upstairs, lobby, warehouse, etc.), a range of related past experiences and potential action associations (or ‘affordances’, to borrow the terminology of Gibson, 1979) are liable to be elicited by the familiar cues now constituting our active sensory flow. Cognitively speaking, the challenge in such situations is not only to recall the appropriate mnemonic episode needed to correctly orient behavior, but also to inhibit any mnemonic associative episodes that threaten to intrude and thereby compete and interfere with retention in conscious attention of a present goal. As many of us know from personal experience, this inhibition of competing interference often fails. The cognitive conflict resolves incorrectly and we begin to perform an action other than our initially intended one. Consequently, we may forget what that original goal was, or perhaps even that we had a goal other than acting upon the intruder content. Only upon arriving back to see our blank notebook do we remember we had gone to the office to get a pen, not this book we’d been thinking about earlier; the pen is still in the bedroom, where we last used it, and so not in the office, despite our clear *prior* memory of using it there.

For an animal performing the EdM test, the central platform of the radial maze is analogous to a lobby of doors, each leading to one of three distinct contexts in which there is a specific, ongoing memory-based task to attend to. Conceiving of the EdM model as analogous to everyday examples such as that above provides an effective lens for bringing into focus the core cognitive processes posited to be at play in such situations: 1) Retrieving a specific episode of mnemonic content (e.g. ‘last location pen was used’ or ‘arm chosen when last in spatial context *A*’) into WM entails inhibition of other environment-related and therefore potentially interfering content (e.g. actions taken in spatial contexts *B* or *C*, or in context *A* but the time prior to last); 2) In turn, this retrieval concomitant inhibition induces a genuine amnesic effect (RIF) upon the inhibited content, making it more difficult to recall at an ulterior stage, perhaps even only moments later. On the flipside, if inhibition of interfering content should fail and another cognitive item successfully intrude into WM, then, following from point 1, this will; 3) Occur concomitantly with a certain level of inhibition of the original voluntarily retrieved content. In which case, finally; 4) This inhibition will induce an amnesic effect upon the original content, creating conditions through which an initial voluntary goal may ultimately be *in*voluntarily over-ruled. Experimentally speaking, potential for mnemonic intrusion during EdM challenge can be further modified via lengthening of the intertrial interval (ITI), being the time the animal is confined to the central platform between each trial. While increase of ITI has classically been understood to increase the demand on memory retention (Al Abed et al., 2016), in the present study we nuance this understanding by emphasizing the increased opportunity for instances of *spontaneous* memory episode replay susceptible to *actively* impact retention during longer ‘offline’ intervals (Xu et al., 2019).

### 1.5 Age-related memory alteration

With respect to physiological effects of ageing on memory function, human studies have shown that organizational cognitive processes are those most often negatively impacted in elderly subjects (for review, see Lustig & Jantz, 2015). In self-report surveys, the kind of ‘mind-wandering’ memory failures illustrated in our ‘forgotten pen’ example are among the most frequently mentioned by elderly subjects (Ossher et al., 2013). Older human subjects are also more likely than young subjects to encode goal-distracting sensory information (Biss et al., 2013), making them more susceptible to elicitation of goal-irrelevant memory items. It has further been shown in both aged humans (Leal & Yassa, 2015) and animals (Gallagher et al., 2010; Wilson et al., 2006) that the formation of new everyday-like memories is hampered by interference from similar pre-existing memories, stemming from the combination of diminished ability to discriminate between similar experiences and overexpression of prior memories (Stark et al., 2010). In this light, concerning the face validity of our model for studying the impact of ageing on organizational memory function, we have previously reported that aged mice display a significant performance deficit under EdM challenge (Al Abed et al., 2016).

### 1.6 Deliberative behaviors

To contribute to a deeper understanding of the cognitive processes underpinning both success and failure in organizational memory function under various physiological and experimental conditions, we further developed novel behavioral analyses for measuring three distinct but interrelated deliberative behaviors elicited by the EdM protocol: 1) Decision latency (time elapsed between trial onset and definitive response choice); 2) Vicarious trial-and-error (VTE, quantification of behavior whereby, at trial onset, an animal begins down one arm but then seemingly ‘changes its mind’ and returns back to the decision point to choose again; first identified and theorized in now classic papers in behavioral psychology; Muenzinger, 1938; Tolman, 1932, 1939, 1948; for review see Redish, 2016); 3) Run-time (time taken for the animal to execute its definitive decision, i.e. travel the distance between the arm threshold and the distal zone of the arm where reward presence or absence will reveal whether the animal has chosen correctly or incorrectly). Our global expectation was that, if RIF does increase as a function of EdM trial difficulty, then a trace of this should be observable not only in the EdM performance profile but also in the deliberation profiles. In concrete terms, if, as laid out by Redish (2016), deliberation is *deliberation over representational contents*, then, in the presence of RIF, we should expect deliberation to counter-intuitively *decrease* as a function of increasing EdM trial difficulty.

### 1.7 Findings

By analyzing the three deliberative behaviors as a function of both trial difficulty and trial outcome (rewarded versus unrewarded), we have been able to demonstrate in the present study that, in young mice, trial difficulty-related increase in EdM errors is accompanied by decrease in deliberation behaviors. By contrast, this trial difficulty-related decrease in deliberation is significantly less pronounced in aged mice, who also engage in significantly more *overall* deliberation than young animals. However, even more strikingly, deliberation behaviors in aged mice are not affected by ultimate trial outcome, unlike in young mice where clear and significant anticipatory traces of outcome can be observed. We interpret these findings as demonstrating that while deliberation requires remembering (i.e. retention of rich representational mnemonic episodes to be deliberated over), anticipation of outcome requires forgetting (e.g. forgetting the prior time we used the pen in the office so we can correctly anticipate finding it in the bedroom, where we actually used it most recently). Regarding the underlying neurobiology of the aged EdM phenotype, we also present results in parallel from a preliminary investigation inspired by recent studies into the role of age-related alteration of the endocannabinoid system in mnemonic decline (Bilkei-Gorzo, 2012; Bilkei-Gorzo et al., 2017). Notably, using the Dlx-CB1-KO transgenic mouse line (Monory et al., 2006), we provide the first evidence of a critical role for cannabinoid type-I receptors expressed on GABAergic neurons of the forebrain (whose signaling is known to alter with age, diminishing or augmenting depending on the area considered: Burjanadze et al., 2022; Ethiraj et al., 2021; Pandya et al., 2019) in successful active forgetting-dependent EdM performance. We observed that Dlx-CB1-KO mice present an extreme version of the aged EdM phenotype while wildtype animals replicate the behavioral profile of young mice. In the discussion section, we refer details of our novel behavioral findings to the cognitive ageing literature into the impact of ageing on memory, on active forgetting, on deliberative decision making, etc. Finally, we propose a tentative unifying neurocognitive interpretation of our results, encompassing a wide range of testable hypotheses and predictions for further future investigation into the processes underlying the novel EdM deliberation profiles described here.

## 2. Materials & Methods

### 2.1 Animals

Young (8-12 weeks) C57BL/6J male mice were obtained from Charles River and collectively housed in a standardized animal room (23 °C; lights on 7 AM to 7 PM; four or five mice per cage). Mice from the aged cohort (20-22 months) underwent ageing in collective housing on site at the animal facility of the Neurocentre Magendie. As previously described (Zerucha et al., 2000; Monory et al., 2006), Dlx5/6-Cre mice were crossed with CB1f/f mice to obtain CB1f/f;Dlx5/6-Cre (here called Dlx-CB1-KO) and CB1f/f (WT) controls. 8-14 weeks old naive male Dlx-CB1-KO and WT animals were used. All animals were moved to individual cages 2 weeks before the beginning of experiments.

#### Food restriction

Five days prior to the first day of training, all animals were placed under a progressive food restriction schedule in order to gradually bring them to 85% to 90% of their baseline free feeding weight. Individual animal weight and welfare was monitored daily throughout the duration of the experimentation. All experiments were conducted in accordance with European Directive 2010-63-EU and with approval from the Bordeaux University Animal Care and Use Committee CCEA50. All efforts were made to minimize suffering and reduce the number of animals used.

### 2.2 Behavior

#### 2.2.1 Apparatus

##### Radial maze

The behavioral apparatuses were 8-armed fully automated radial mazes (Imetronic). The surface of each maze is raised ∼100cm off ground level. Access to each arm is from a central platform by means of automated vertically retracting doors. When all doors are closed during behavior, the animal is contained within the central platform, a regular octagon of size ∼485cm² and edge 10cm (i.e. the width of each arm and door). At the distal end of each 50cm length arm is an automated pellet distributor for dispensing food reward. The distributor is set into a slight indent in order to hide its state (i.e. baited or not baited) from the animal. Animal movements in the radial maze are detected via video camera and motion detection software (GenCam). The motion detection software communicates with a second piece of software, POLYRadial (Imetronic), through which pre-programmed sequences of automated radial maze actions are triggered. This program is used for the design and execution of behavioral exercises (sequences of door openings, location of food reward, conditions for opening and closing of doors, etc.). Hence, the exercises are customizable and contingent upon a combination of both the detected movements of the animal and automated timed sequences.

#### 2.2.2 Behavioral model

##### Habituation

Prior to the first day of everyday-like memory (EdM) testing, all animals were habituated to the context and functioning of the radial maze apparatus. Food restriction, as described above, began three days before habituation (i.e. five days before training). At the beginning of each habituation session, the animal was placed by the experimenter in the central platform of the radial maze, all 8 doors of which were closed. Once removed to the control room, the experimenter launched the habituation program via the POLYRadial software. The habituation program began by an interval of 10 seconds during which the animal could explore the central platform. Following this, all 8 doors opened simultaneously, presenting the animal with the opportunity to freely explore the entire surface of the maze. As the animal explored, once it had advanced to the most distal section of a given arm (location of the distributor and food reward) and returned to the central platform, the door of that arm automatically closed behind it, preventing further access to that arm in the current session. Thus, once the animal had fully explored all 8-arms, it found itself again contained within the central platform. At this point, a further habituation session could be launched if needed. It was considered that when an animal had recovered and consumed at least 5 out of 8 available food rewards in a single session that it was fully habituated to the relevant functionalities of the apparatus.

##### EdM protocol

As previously described (Al Abed et al., 2016), the aim of the EdM radial maze protocol is to model daily life situations in which numerous, repetitive events, varying in temporal distance and often with only subtle but crucial changes in their contextual content, must be accurately encoded in memory for later context-appropriate retrieval. Demands on mnemonic retention versus selective organization are inversely proportional with respect to the temporal interval between context repetitions. By way of illustration, the more frequently I park my car in the same zone, the lower the demand on mnemonic retention (e.g. “Where did I park at work this morning?” implies less demand on retention than “Where did I park at the airport two weeks ago?”), but the higher the demand on inhibitory mnemonic organization in order not to mix up successive placements provoking ‘proactive interference’ (“This isn’t my car! This is where I parked *yesterday* morning.”).

Responding to the EdM challenge (figure 1A), animals must simultaneously (i) mnemonically encode spatiotemporally relevant items of information regarding what actions they are taking on three distinct task contexts (arm pairs *A, B*, and *C*), and (ii) use and update each of these mnemonic items in order to choose the correct action on subsequent trials (according to a simple rule of ‘always choose the action not chosen during the previous trial on a given pair’). In other words, in the course of each trial, they must memorize which arm (spatial) they have chosen that time (temporal) and, on the following trial in the same task context, choose the other arm (figure 1B). Identification of which pair is which depends on spatially identifying its position relative to prominent extra-maze cues in the experimental room (represented by the black shapes surrounding the maze in figure 1A). Mice have no way of predicting which of the three task contexts they will be presented with on any given trial and thus no way of knowing, while waiting in the central platform for the next trial to begin, which memory episode they will next need to retrieve nor, likewise, which ones they will need to inhibit.

In each session of EdM, animals are presented with a sequence of 23 trials, with each trial consisting of one presentation of one of the 3 arm pairs (*A, B*, and *C*). This sequence changes from session to session and is pseudo-random (i.e. unpredictable) from the subject’s perspective. On each trial, mice must choose to visit one arm out of a presented pair of arms (always neighboring). The food reward will always be located at the distal end of that arm which was not visited by the mouse on the previous presentation of the same task context, independently of whether that previous choice was correct or incorrect. In other words, the reward in a given task context switches location only once it has been retrieved (compare what happens with pairs *A* and *C* to what happens with pair *B* in the sample sequence represented in figure 1B). The task therefore implies mnemonically encoding and storing information relative to which arm was visited on any given arm pair presentation n-1 until the next trial n consisting of a presentation of the same arm pair. In short, the EdM protocol relies on and also reinforces, according to a ‘win-shift’ strategy (McDonald & White, 1993), the spontaneous mouse behavior of spatial alternation. The number of interposed trials on either of the other two task contexts, plus the duration of the inter-trial intervals (ITI), together constitute the retention component of the task. However, retention in the context of the EdM protocol is highly dynamic and, for any given trial, a significant organizational mnemonic component is present in both the necessity to inhibit spatially irrelevant content pertaining to the *n-1* trials on the other two pairs as well as the need to inhibit spatially relevant but temporally (or episodically) irrelevant pair-specific interference on any given arm pair presentation *n*. This is to ensure that the prior *n-1* choice and not the prior-prior *n-2* choice on the present pair is acted upon (figure 1B). Both these varieties of trial *n* context irrelevant memory episodes constitute interfering and competing cognitive noise in the decision making process.

At the beginning of each EdM session, the animal is placed at the center of the maze, with all vertically opening arm access doors in the up/closed position. After a short pause, the two doors to one of the 3 pairs (*A, B* or *C*) open simultaneously, retracting vertically below the surface of the maze, whereupon the mouse can enter either of the two neighboring arms. Only once the mouse has reached the distal reward zone of what at that instant becomes its definitively chosen arm does the door to the non-chosen arm close, preventing further revision of choice. When the mouse returns from the distal zone and is again detected in the central platform of the maze, the door of the chosen arm also closes behind it and the mouse finds itself again fully enclosed in the central platform. After a given ITI (in the present study ITIs <= 10s or = 30s), during which the mouse is confined to the central platform, another trial begins with the opening of two doors, either those of the same task context (arm pair) again or those of one of the other two task contexts, and so on. During the initial trial on each pair per session, both arms contain a food reward. These three initial ‘sampling’ trials thereby serve to establish how the reward will spatially alternate in all subsequent trials and so are not included in the calculation of EdM performance. So, out of 23 total trials per session spread across 3 pairs, EdM performance is calculated from a subject’s mnemonic decision making behavior on 20 of those trials.

The mnemonic challenge of EdM upon which we primarily focus our analyses in the present study is determined by the number of trials in the other two task contexts interposed between a presentation *n-1* and subsequent presentation *n* of one and the same task context. This gives rise to 5 levels of difficulty, denoted by the number of such interposed trials, i.e. from 0 to 4. Level 0 therefore corresponds to two immediately consecutive presentations of the same arm pair, with no interposed trials on the other pairs. Level 0 thereby most closely resembles a classical T- or Y-maze spontaneous alternation trial (figure 1B; the second presentation of pair *C* is a level 0 trial; the second presentation of pair *A*, a level 1 trial; the third presentation of pair *A*, a level 3 trial, etc.). However, in crucial contrast to a classical T- or Y-maze, even on level 0 EdM trials on a given pair, there is still a matter of environmentally related but task context irrelevant mnemonic competition from memories of actions taken on the other two pairs.

The sequences of pairs *A, B*, and *C* within a given training session of the EdM model have been designed such that over three consecutive sessions the number of trials of each difficulty level are equally balanced. Hence, behavioral parameters expressed as a function of trial difficulty are calculated as either the mean or median per block of 3 sessions and analyzed on this basis. Each block of 3 sessions consists of 12 trials at each complexity level, for a total of 60 trials, plus the 3×3 initial ‘sampling’ trials in each session which do not figure in evaluation of the EdM performance. Results presented here represent the merged mean or median of between 2 and 4 such training blocks.

#### 2.2.3 Deliberative behaviors

##### Decision latency

Time elapsed between the onset of a trial (doors of the current trial pair open) and the instant when the threshold from the central platform into the surface-arm of the animal’s definitive choice is first crossed (decision latency, milliseconds).

##### Vicarious trial-and-error/VTE

During certain trials, animals may either entirely or almost entirely cross the threshold into one arm of a pair but, rather than continuing to the distal reward zone (which would trigger the door to the non-chosen arm to close), instead physically retreat out of the arm. At this point, animals may either choose to explore the other arm of the pair or (more rarely) re-enter the initially ‘chosen’ arm. As long as the distal zone of either arm has not been entered, this process can technically continue indefinitely. We interpreted this behavior as an occasional overt and full body manifestation of the ongoing ‘covert’ cognitive deliberation process. On a given trial, each additional crossing of either of the two central platform-to-arm thresholds, in the direction from the platform towards the arm only, was quantified as one unit of VTE. We refer to these as ‘KOOK’ units, a label intended to capture the fact that VTE choice revisions are ultimately either error-inducing (‘KO’) or rectifying (‘OK’).

##### Technical side-note on VTE

This behavioral phenomenon was first identified and theorized in now classic papers in behavioral psychology (Muenzinger, 1938; Tolman, 1932, 1939, 1948; Hu & Amsel, 1995; Bett et al., 2012; Huynh et al., 2021; for review see Redish, 2016). There are, however, some important differences between this classical VTE and what we report here under the same label. The VTE that has most often been studied in rats is ethologically more fine-grained, since it quantifies not only full-body movements but also side-to-side head movements resulting from an animal looking back and forth between options. Though we did observe our mice making such head movements, the scale of our radial maze apparatus compared to the body size of a mouse meant that the sensitivity of our gridline motion tracking equipment allowed us to identify and quantify only large or full-body movements. We therefore suggest that the large movement behavior we report here as VTE is only a representative overt fraction of the full range of cognitive VTE behavior occurring both internally (i.e. ‘covertly’ or ‘mentally’) and in a range of discrete to large physical movements. Redish (2016) has already suggested something similar even in the case of rats, thus our claim is simply that this is all the more true in our mouse study. Despite its relatively more coarse-grained nature when applied to mice rather than rats, the VTE behavior nevertheless constitutes robust and fertile ground for further, deeper investigation into the underlying cognitive and neural processes of EdM deliberation: during the primary experiments presented here, 41 animals performed a total of 1,147 identifiable VTE events over a total of 12 sessions.

##### Run-time

Time taken to execute the definitive decision, i.e. travel the distance from the threshold of the chosen arm to the reward-distributor containing distal zone (run-time, milliseconds). We interpret this behavioral parameter as a proxy measure for response vigor related to choice confidence/hesitancy.

#### 2.2.4 Analysis

##### Code

All raw data extraction, analysis, statistical comparison, and graphical representations were generated using custom codes written in Python (Van Rossum & Drake, 2009) using the pandas (Reback et al., 2020), numpy (Harris et al., 2020), pingouin (Vallat, 2018), bioinfokit (Bedre, 2021), matplotlib (Hunter, 2007), and seaborn (Waskom, 2021) libraries. All code is open source and available either upon request or at https://github.com/metaphysiology.

##### Statistical analyses

Following the previous relevant literature (Marighetto et al., 2011; Al Abed et al., 2016), EdM performance data was analyzed using ANOVA with post-hoc tests. Similar ANOVA with post-hoc analysis was applied to the VTE data, which was quantified and graphically displayed as mean population values. Global ANOVA analyses included between-sample factors such as ‘Group’ (Young, Aged) or ‘ITI’ (10s, 30s) and within-subject factors such as ‘Difficulty’ (levels 0-4) or ‘Outcome’ (Correct, Incorrect). Temporal decision latency and run-time measures were quantified and graphically displayed as median population values and were therefore statistically analyzed as nonparametric data sets using nonparametric equivalents to ANOVA (i.e. Kruskal-Wallis test or Friedman analysis for within-subject repeated measures) with suitable post-hoc tests (i.e. Wilcoxon, Mann-Whitney U test). Significance level was set to *p* < 0.05 and all graphical representations are similarly displayed with an error band marking the bounds of a 95% confidence interval.

## 3. Results

### 3.1 Everyday-like memory performance

#### 3.1.1 EdM errors increase as a function of trial difficulty

Operationally speaking, the difficulty level of any trial *n* in a given pair-context equates to the total number of trials in the other two pair-contexts interposed between the prior trial *n-1* in the same context and *n* itself. For example, a trial *n* of difficulty level 3 on pair *A* entails that, since completion of trial *A*_*n-1*_, three interposed trials on pairs *B* and/or *C* have since been executed (figure 1). With respect to a given *A*_*n-1*_ memory episode, a trial *A*_*n*_ of difficulty level 3 thereby represents on average three times more occasions than a trial of difficulty level 1 for *A*_*n-1*_ to have arrived into the representational WM deliberation space and consequently been ‘actively forgotten’ as interfering content (i.e. as irrelevant to trials in spatial contexts *B* and *C*) prior to being required and actively retrieved at the onset of trial *A*_*n*_. Our first objective was therefore to replicate previous confirmation by our team (Al Abed et al., 2016) of the consequent hypothesis that significantly more EdM errors will occur on higher compared to lower difficulty trials. The novelty of the present study is the further hypothesis that greater levels of RIF elicited by higher difficulty trials is the neurocognitive mechanism underpinning the baseline (i.e. as observed in young healthy mice at low ITI) EdM trial difficulty performance profile (figure 2, left, red curve, and; supplementary figure S.1A, red curve).

**Figure 2.**
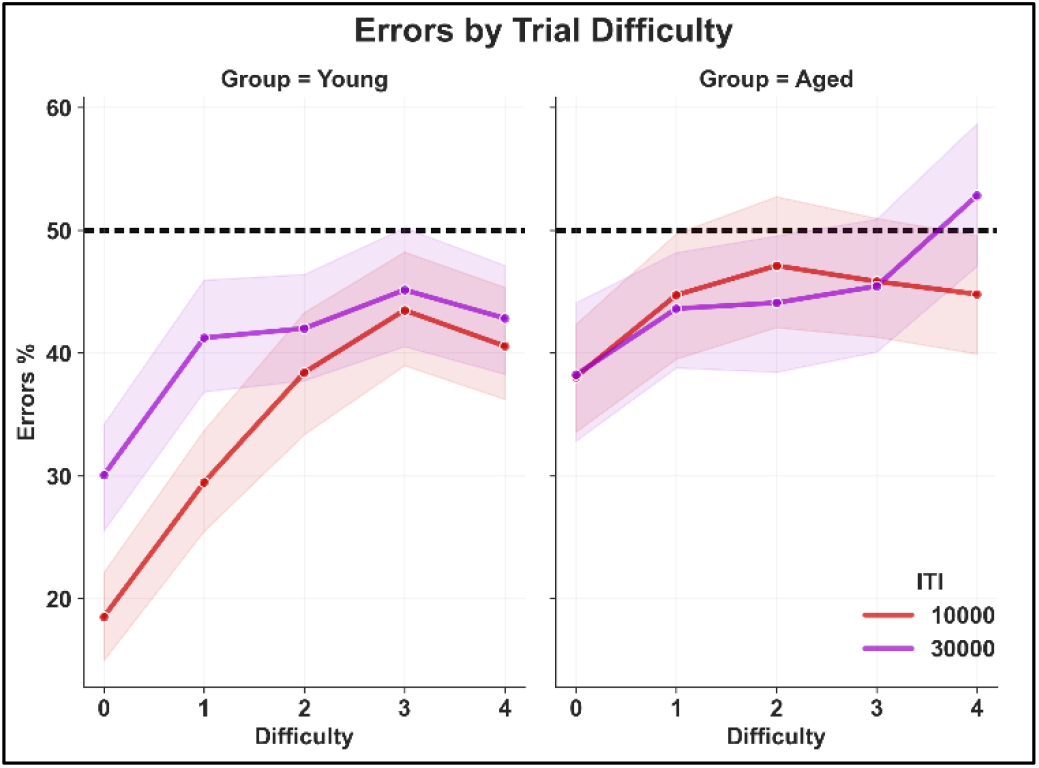
Everyday-like memory errors increase as a function of trial difficulty, but impact of trial difficulty, ITI, and ITI*difficulty differs in young versus aged mice. Experimental groups (‘ITIGroup’ in analyses): young ITI 10s (left column, red curve; *n* = 16); young ITI 30s (left column, indigo curve; *n* = 16); aged ITI 10s (right column, red curve; *n* = 8); aged ITI 30s (right column, indigo curve; *n* = 8). For certain analyses, these groups were pooled by age into Young and Aged (‘Group’ in analyses and figures). Results are averaged over multiple EdM training blocks, giving 12 to 48 trials per trial difficulty per animal. All error bands represent 95% confidence intervals; vertical spaces between bands provide visual indication of statistical significance. Horizontal line at 50% represents level of errors expected when decision making is random. Initial three-way ANOVA revealed significant main effects of ‘Group’ (i.e. age) ‘ITI’, and ‘Difficulty’, plus significant interactions of ‘Group*ITI’ and ‘Group*Difficulty’. Post-hoc pairwise Tukey tests revealed that the ‘Group*ITI’ effect was primarily driven by the ‘Young*ITI10s’ population. Within the ‘Young*ITI10s’ population, post-hoc pairwise t-tests revealed a significant effect of ‘Difficulty’ between trial difficulty levels 0 and 1 and respective higher levels, but in the ‘Young*ITI30s’ and ‘Aged*ITI10s’ and ‘Aged*ITI30s’ populations, a significant impact of trial difficulty was observed only between level 0 and higher levels but not between any of the other levels (e.g. 1 vs 3 or 1 vs 4, etc.). Moreover, in aged mice, this significant trial difficulty effect was only observable when higher levels were combined and compared as a block, using T-tests, against level 0 error values. See main text for details of statistical analysis.

An initial three-way ANOVA analysis of the data (figure 2) revealed significant main effects of ‘Group’ (i.e. age), ‘ITI’, and ‘Difficulty’ (*F*_(1, 2218)_ = 44, *p* < 0.0001; *F*_(1, 2218)_ = 12.6, *p* < 0.0001; *F*_(4, 2218)_ = 25, *p* < 0.0001, respectively), plus significant interactions of ‘Group*ITI’ (*F*_(1, 2218)_ = 6, *p* = 0.01), ‘Group*Difficulty’ (*F*_(4, 2218)_ = 3.5, *p* = 0.007), and ‘Group*ITI*Difficulty’ (*F*_(4, 2218)_ = 2.4, *p* = 0.05). To directly test the hypothesis that more errors occurred on higher difficulty trials, we performed one-tailed post-hoc independent T-tests with Bonferroni correction and repeated measures (within ‘Difficulty’). Beginning with the ‘Young*ITI10s’ population as a baseline (figure 2: left, red curve), these tests revealed that significantly more errors occurred at difficulty levels 1 – 4 relative to level 0 (*p*-values <= 0.002) and that this effect ramped up in size across levels 1 – 3 before flattening out/slightly decreasing again at level 4 (abs. Cohen effect sizes; *d* = 1.3, 2, 2.5, 2.1, respectively).

Relative to level 1 also, a nearly statistically significant trend (level 3 reaching *p* = 0.06 after correction) for more errors to occur on higher difficulty trials (levels 2 – 4) was observed, with the same ramping then flattening out of effect sizes (abs. Cohen effect sizes; *d* = 0.6, 0.9, 0.7). Indeed, in the ‘Young*ITI10s’ population, we observed a general trend for more errors to occur at all higher relative to lower trial difficulties, with the exception of level 4 where slightly but not significantly fewer errors occurred relative to level 3.

#### 3.1.2 Increased ITI and age associated with increase in errors and shift towards more ‘binary’ performance slope

Post-hoc pairwise Tukey tests revealed that the significant ‘Group*ITI’ effect was driven primarily by all other ‘Group*ITI’ populations committing significantly more overall errors compared to the baseline ‘Young*ITI10s’ population (*p*-values <= 0.0002). Only one other significant difference between populations was observed in overall error numbers, that between the ‘Aged*ITI30s’ and‘Young*ITI30s’ populations (*p* = 0.02), though the effect size of this difference was smaller than that between the ‘Young*ITI30s’ and ‘Young*ITI10s’ populations (abs. Cohen effect sizes, *d* = 0.17 vs *d* = 0.23, respectively). In other words, the ‘Young*ITI30s’ population displayed more closely comparable numbers of EdM errors to the ‘Aged*ITI10s’ and ‘Aged*ITI30s’ populations than to the ‘Young*ITI10s’ population. In short, compared to baseline experimental conditions, increase of either ITI or age significantly increased overall number of consequent EdM errors.

With respect to the ‘Group*ITI*Difficulty’ interaction, in the ‘Young*ITI30s’ population (figure 2, left, indigo curve), significantly more errors were again observed at higher difficulty levels relative to level 0 (lvs. 1 – 4, *p*-values with Bonferroni correction <= 0.008), although with noticeably less ramping in the successive effect sizes when compared to the ‘Young*ITI10s’ population (abs. Cohen effect sizes; *d* = 1.1, 1.2, 1.2, 1.2, respectively). Indeed, no significant difference in error numbers was observed in the ‘Young*ITI30s’ animals between level 1 and any higher difficulty levels. In the ‘Aged*ITI10s’ and ‘Aged*ITI30s’ populations, we observed only a very slight and non-significant trend for more errors to occur at each of the higher difficulty levels relative to level 0. Almost no difference was observed in error values between any of the other trial difficulty levels in either of the Aged*ITI populations. Hence, EdM performance in aged mice contrasted with that observed in young mice in that no statistically significant differences in mean errors were observed between any of the individual difficulty levels, certainly related to aged mice committing more errors even at level 0 than young mice.

However, since we posit that trials of difficulty level 1 or higher require episodic memory (EM) function while level 0 trials *can* be successfully accomplished using only the retention dimension of working memory (WM), this presents two qualitatively, as opposed to merely quantitatively, distinct categories of mnemonic EdM challenge. Hence, we also performed one-tailed T-tests which revealed significantly lower mean error values on level 0 trials relative to the mean error values of all higher levels combined (‘Aged*ITI10s’, *t*_(155)_ = -2.8, *p* = 0.003; ‘Aged*ITI30s’, *t*_(141)_ = -2.6, *p* = 0.005). In this sense, a similarity between the ‘Young*ITI30s’ and two Aged*ITI populations was observed: although varying in significance, all three displayed a ramp in errors between levels 0 and 1 followed by a relative levelling out of error numbers between levels 1 to 4, the slight but not significant increase in errors at level 4 in the ‘Aged*ITI30s’ population only notwithstanding (figure 2, right, indigo curve). Whether this latter discrete increase only on difficulty level 4 trials bears a meaningful relationship to the increase in ITI or is simply a result of random performance fluctuations is a question that will require further replication and investigation.

#### 3.1.3 Aged mice performance not impacted by ITI

In stark contrast to young mice, increased ITI appeared to have little meaningful impact on EdM performance in aged mice, as corroborated by a TOST equivalence test which showed that the proportion of errors in the ‘Aged*ITI10s’ versus ‘Aged*ITI30s’ populations showed significant equivalence to within a bound of 4% (*p* = 0.03).

To summarize the EdM performance results; as trial difficulty, ITI, and age increased, the mean proportion of error values tended to increase also, approaching an asymptotic high error plateau close to random performance levels (i.e. approaching 50% errors), such that on trials of difficulty 3 – 4 almost no differences in EdM performance could be observed between any of the populations, either as a function of age or ITI. It is, however, worth noting that in the ‘Young*ITI10s’ population only, although error values on the most difficult trials approached random performance levels, they still remained significantly below this line across all difficulty levels (figure 2, left; see 95% confidence interval error bands of red line vs 50% line). In the ‘Young*ITI30s’, ‘Aged*ITI10s’, and ‘Aged*ITI30s’ groups, the ‘plateau’ of non-significant difference spanned all trial difficulties greater than 0, i.e. across all putatively EM- (as opposed to WM-) dependent EdM challenges. Across all 4 populations, the greatest performance differences were observed between level 0 and all other levels. Up to this point, the performance results we observed replicate the central elements of similar EdM studies previously published by our lab (Al Abed et al., 2016).

#### 3.1.4 Preliminary results into neurobiological basis of EdM deficit

With respect to our hypothesis that the relative smoothing of the EdM performance profile as a function of age was due to an age-related active forgetting deficit, and given that age-related mnemonic decline has been linked to age-related decreases in endocannabinoid system (eCS) function (Bilkei-Gorzo, 2012; Bilkei-Gorzo et al., 2017), we also analyzed EdM data from parallel experiments using (young) mice in whom cannabinoid type-I receptors (CB1R) had been deleted from GABAergic neurons of the forebrain (Dlx-CB1-KO mouse line; see Materials & Methods). Interestingly, given that the literature has demonstrated that this mouse line displays no deficit in simple T- or Y-maze spatial alternation working memory tasks (Albayram et al., 2016), we nevertheless observed that Dlx-CB1-KO animals performing at ITI values of 10s or lower were even more impaired than aged mice in the specifically active forgetting-dependent EdM challenge, barely performing fewer errors than could be explained by random choice behavior even on level 0 trials (supplementary figure S.1A, green curve). By contrast, wildtype animals from the same experiments almost identically replicated the EdM performance profile of our baseline ‘Young*ITI10s’ population (supplementary figure S.1A, red curve).

### 3.2 Deliberation behaviors

We next turned to analysis of the three deliberative behaviors (decision latency, VTE, and run-time; see Materials & Methods for details) in order to test our hypothesis that, due to the increased RIF entailed by higher difficulty trials, these behaviors should decrease as a function of increasing EdM trial difficulty. To be tested also was the other side of the same hypothesis, i.e. that due to putative age-related reduction in RIF, we would see an increase of deliberation in aged mice. Given frequent inter- and intra-individual outlier values, we analyze and graphically display the temporal deliberative behaviors (decision latency and run-time) as non-parametric distributions, using the median rather than the mean population values throughout.

#### 3.2.1 Deliberation decreases as a function of increasing trial difficulty

In initial analyses across all animals, repeated measures Friedman or ANOVA tests revealed highly significant effects of trial difficulty on decision latency (figure 3A; *F*_(4, 186)_ = 15, *p* < 0.0001), VTE (figure 3B; *F*_(4, 188)_ = 5.3, *p* = 0.006), and run-time (figure 3C; *F*_(4, 186)_ = 4.6, *p* = 0.001). In all three, one-tailed pairwise post-hoc Wilcoxon or Tukey tests revealed that, as hypothesized, the trend of this effect was towards significantly less deliberation on higher difficulty compared to lower difficulty trials, with the largest effect sizes observed when comparing between level 0 and higher level trials. However, further repeated measures Friedman or ANOVA tests within each ‘Group*ITI’ population revealed important and varied differences at this level. Notably: the effect of trial difficulty on decision latency lost significance at ITI30s in aged mice; trial difficulty had no significant effect on overall VTE in aged mice at either ITI value or in the ‘Young*ITI10s’ population (though a *p* = 0.08 trend was observed here and is lent further weight by trial difficulty having a significant effect on VTE in the WT population from the Dlx experiments; supplementary figure S.1C, *F*_(4, 74)_ = 25, *p* < 0.0001); and the effect of trial difficulty on overall run-time did not reach significance in either the ‘Young*ITI10s’ or ‘Aged*ITI30s’ populations. [Note however that a deeper underlying significance of the impact of trial difficulty on overall deliberative behaviors will be brought to light when further analyzed as a function of trial outcome (see below, figure 4 and related main text).]

**Figure 3.**
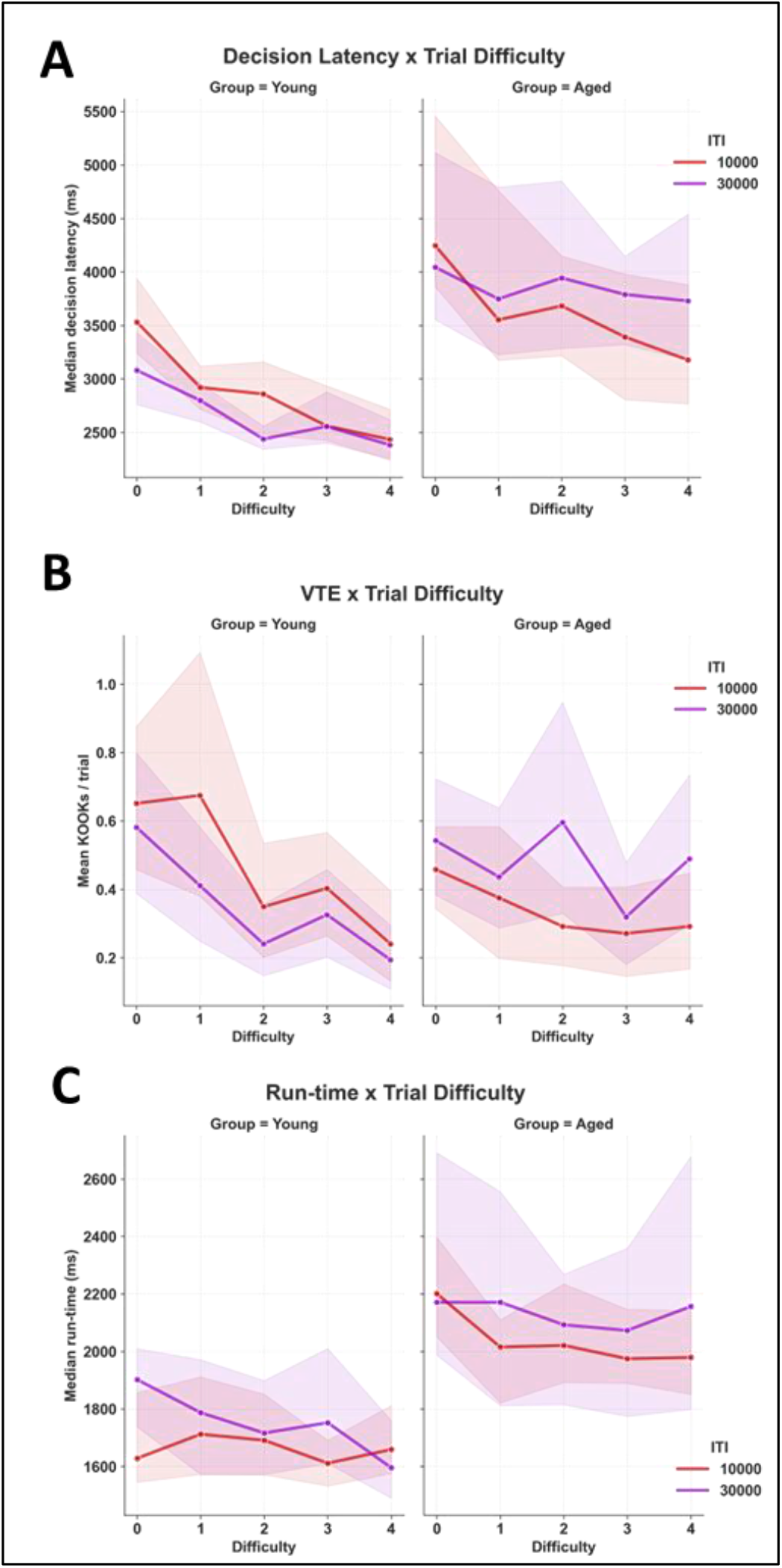
Deliberation behavior increases with age and increasing ITI produces opposing effects in young vs in aged mice. Experimental groups: ‘Young*ITI10s’ (left columns, red curves; *n* = 16); ‘Young*ITI30s’ (left columns, indigo curves; *n* = 16); ‘Aged*ITI10s’ (right columns, red curves; *n* = 8); ‘Aged*ITI30s’ (right columns, indigo curves; *n* = 8). For certain analyses, these groups were pooled by age into Young and Aged (‘Group’ in analyses and figures). Temporal results (decision latency and run-time) are displayed as median population values, VTE as population means. Results pooled across multiple EdM training blocks, giving 12 to 48 trials per trial difficulty per animal. All error bands represent 95% confidence intervals; vertical spaces between bands provide visual indication of statistical significance. **A & C)** Independently of ITI, age had a significant effect on both decision latency and run-time. **A–C)** Averaged across all Group*ITI populations, a highly significant effect of difficulty was observed on decision latency, VTE, and run-time, whereby less deliberation was observed on trials of higher compared to lower difficulty. Post-hoc tests revealed this trial difficulty effect was primarily but not exclusively driven by the young populations. **A–B)** In the two pre-definitive choice deliberative behaviors (i.e. decision latency and VTE) ITI was observed to have opposing effects on young vs aged mice, decreasing deliberation in the former but increasing it in the latter. A bootstrapped resampling approach (to compensate for the unpaired population study design) revealed this divergence of effects to be statistically significant (see main text for details).

**Figure 4.**
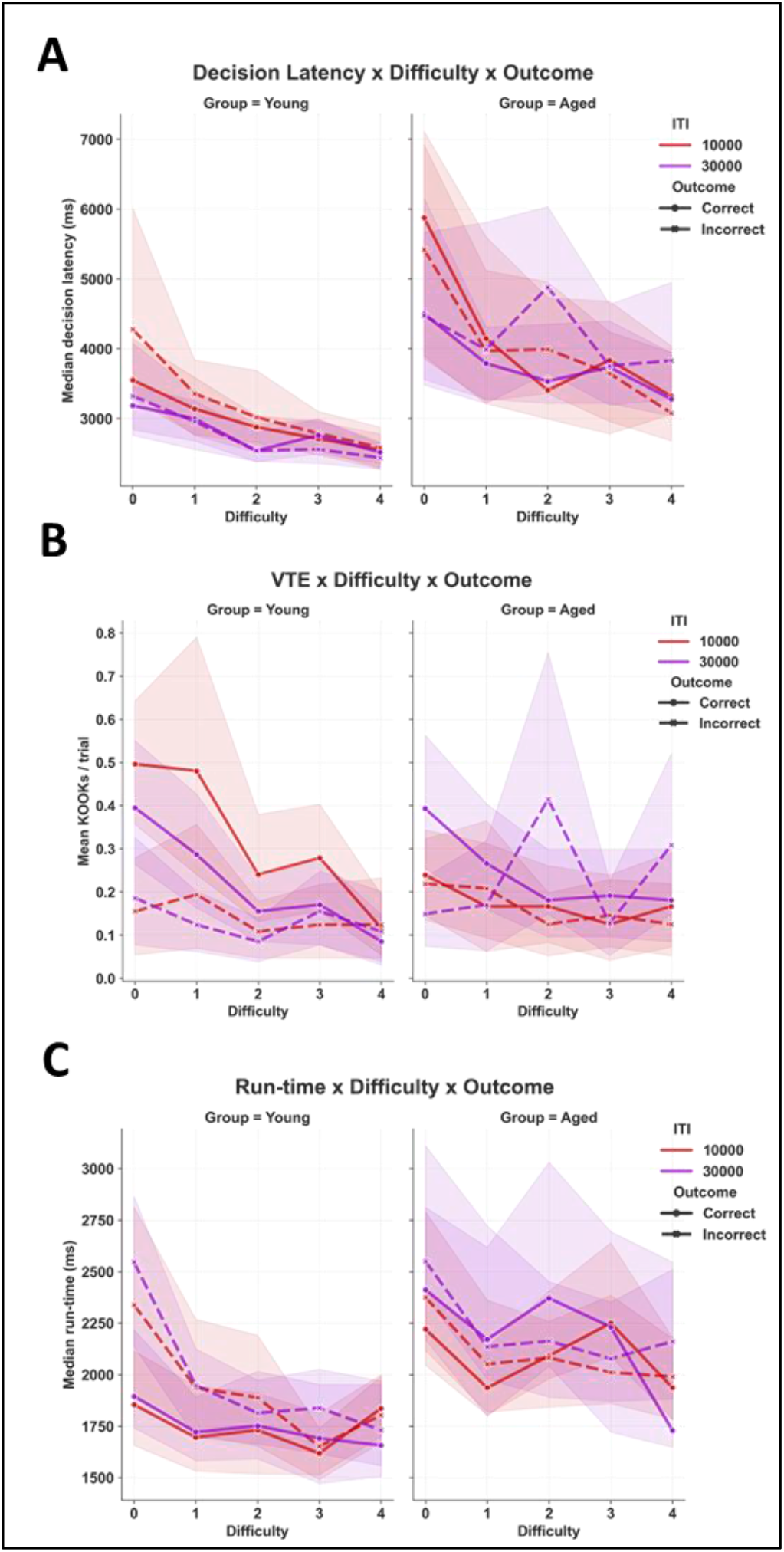
Anticipatory cognition leaves significant behavioral traces in deliberation in young but not in aged mice. Experimental populations: ‘Young*ITI10s’ (left columns, red curves; *n* = 16); ‘Young*ITI30s’ (left columns, indigo curves; *n* = 16); ‘Aged*ITI10s’ (right columns, red curves; *n* = 8); ‘Aged*ITI30s’ (right columns, indigo curves; *n* = 8). For certain analyses, these populations were pooled by age into Young and Aged (‘Group’ in analyses and figures). Temporal results (decision latency and run-time) are displayed as median population values, VTE KOOK units as population means. Results pooled across multiple EdM training blocks, giving 12 to 48 trials per trial difficulty per animal. All error bands represent 95% confidence intervals; vertical spaces between bands provide visual indication of statistical significance. **A)** An apparent trend for longer decision latency prior to incorrect choices was seen in the ‘Young*ITI10s’ population only. Although not statistically significant here (repeated measures Friedman within ‘Outcome’, *p* = 0.14), the same trend was observed to a significant level in WT animals from the Dlx-CB1-KO experiment [supplementary figure S.1B]. No anticipatory differentiation of decision latency was observed in either the aged or ‘Young*ITI30s’ populations. **B)** A significant impact of ‘Outcome’ was observed in VTE in young but not in aged mice. In young animals, the significant decrease of VTE as a function of ‘Difficulty’ was revealed as a property only of choice rectifying VTE. **C)** A significant impact of ‘Outcome’ was similarly observed in run-time in young but not in aged mice. In young animals, decrease of run-time as a function of ‘Difficulty’ was revealed as a property only of the run-time of ultimately unrewarded trials.

With respect to the relation between trial difficulty and deliberation, in the case of decision latency and VTE, our observations had led us to hypothesize that at least part of the reason these values were often so much significantly higher on level 0 trials (recall, level 0 = directly successive presentations of the same pair with no interposed trials on any other pair; see Materials & Methods) could be due to what in humans is referred to as ‘rehearsal’ behavior (Basile, 2018). Such behavior consists in the cognitive and often physica repetition of an episode which a subject predicts will be required for recall at a later stage, such as vocally repeating a new phone number upon hearing it. In short, if in the course of EdM challenge, trials are indeed ‘rehearsed’ upon completion, then we might expect this process to temporally and/or physically encroach into the decision latency and/or VTE windows of immediately subsequent level 0 trials, especially at lower ITI values. However, strictly speaking such ‘rehearsal’ would not fall under the category of a deliberative process (see ‘Rehearsal’ section of Discussion for further details). Hence, in light of this possibility, we considered it important to verify whether trial difficulty still had a significant effect on decision latency and VTE even if level 0 trial behaviors were excluded from analysis. In the case of decision latency under these conditions, only the ‘Young*ITI10s’ population retained an impact of trial difficulty that approached statistical significance (Friedman repeated measures within ‘Difficulty’, levels 1-4, *F*_(3, 43)_ = 2.7, *p* = 0.06), though this result was significantly replicated in the WT population from the Dlx-CB1-KO experiments (*F*_(3, 55)_ = 6.2, *p* = 0.001). In VTE, the impact of trial difficulty was maintained only as an almost significant trend in the ‘Young*ITI30s’ population (repeated measures ANOVA with Greenhouse-Geisser correction, levels 1-4, *F*_(3, 45)_ = 2.9, *p* = 0.07). Again, however, the same trend was found to be highly significant in the WT population from the Dlx-CB1-KO experiments (*F*_(3, 57)_ = 6.8, *p* = 0.002).

#### 3.2.2 Decision latency and run-time increase with age

Our graphic representations (figure 3) also pointed towards important differences in deliberative behaviors as a function of age, corresponding to hypotheses formulated on the basis of putatively reduced RIF in aged animals. Confirming this, Kruskal-Wallis or ANOVA comparisons with two-tailed pairwise post-hoc Wilcoxon or Tukey tests revealed a highly significant effect of age on decision latency (figure 3A; between ‘Group’; *H* = 146, *p* < 0.0001) and run-time (figure 3C; *H* = 116, *p* < 0.0001), but not on VTE (figure 3B; *F*_(1, 2238)_ < 0.01, *p* = 0.99). However, comparing VTE behavior between young and aged populations at ITI30s we did observe a more significant effect of age (*p* = 0.03) than that observed between young and aged populations at ITI10s (*p* = 0.05), an observation which demanded further investigation.

#### 3.2.3 Subtle age-differentiated impact of ITI on deliberation behaviors

In contrast to the clear impact of age on deliberative behaviors, the possibility of a more subtle and complex effect of ITI and its interaction with age presented itself. First, in decision latency (figure 3A), an initial one-way Kruskal-Wallis analysis across all populations showed no main effect of ITI (*H* = 0.4, *p* = 0.5). However, a subsequent two-way ‘Group*ITI’ pairwise Mann-Whitney U test (an independent T-test for non-parametric data) with Welch correction confirmed that the decreased decision latency apparent in the ‘Young*ITI30s’ population was statistically significant (*U* = 223075.5, *p* = 0.04). By contrast, the converse apparent increase in decision latency apparent in the ‘Aged*ITI30s’ was not in itself statistically significant (*U* = 107103.0, *p* = 0.36). Similarly, in VTE (figure 3B), a two-way ANOVA showed no main effect of either ‘Group’ or ‘ITI’ but did reveal a significant interaction effect of ‘Group*ITI’ (*F*_(1, 2236)_ = 8.5, *p* = 0.004). However, a further Tukey post-hoc test found no significant difference in VTE between ITI values within either the young or aged mice, despite trends apparent in the graphical representations of the data.

Based on the above results and trends, and on our hypothesis of contrasting RIF functionality in aged versus young mice, we wished to further explore whether increase of ITI quantifiably drove pre-definitive choice deliberative behaviors (decision latency and VTE but not run-time) in opposing directions according to age; i.e. towards less deliberation in young mice but more deliberation in aged mice. In order to test the strength of this hypothesis in the most robust manner possible, given that our data was drawn upon independent populations, we employed a bootstrapping approach whereby the ‘Young*ITI10s’ vs ‘Young*ITI30s’ and ‘Aged*ITI10s’ vs ‘Aged*ITI30s’ data was iteratively resampled (1000 iterations) to create random within-group pairs from which, at each iteration, the delta differences between decision latency and VTE at ITI10s versus ITI30s could be estimated. A subsequent Mann-Whitney U test on decision latency and two-tailed T-test on VTE revealed that the differences between the subsequent aggregated [‘Young*ITI10s’ – ‘Young*ITI30s’] versus [‘Aged*ITI10s’ – ‘Aged*ITI30s’] data were highly significant (figure 3A; decision latency, *U* = 183908, *p* < 0.0001: figure 3B; VTE, *t*_(1058)_ = 4.2, *p* < 0.0001). While such a bootstrapped result calls for further behavioral replication, it reinforces the basis for the visually apparent trend that increasing ITI does produce significantly contrasting cognitive effects in young versus aged mice (figure 3A-3B). Why this might be the case with respect to the two pre-definitive choice deliberation behaviors (i.e. decision latency and VTE) but not the post-definitive choice one (i.e. run-time) will be broached in the discussion section.

#### 3.2.4 Strategic value of VTE behavior

Since VTE only occurs on a subsample of trials, we next analyzed EdM performance as a function of occurrence of VTE behavior in order to determine whether or not engaging in VTE was a globally successful cognitive strategy. To do this, we calculated per trial performance as a function of whether either 0 or more than 0 KOOK units of VTE had occurred (1 KOOK unit = one crossing of arm threshold with subsequent retreat back to centre; see Materials & Methods). One-tailed T-tests on this data revealed that in young mice only, engaging in VTE was correlated with fewer mean errors than not engaging in it, demonstrating that VTE represented a successful cognitive strategy (*t*_(562)_ = 1.7, *p* = 0.04), albeit a strategy whose cognitive ‘availability’ seemingly decreased as a function of trial difficulty/putative repeated active forgetting. In aged mice, by contrast, whether VTE was engaged in or not had no observable impact on performance, indicating there was no strategic reason for them to either engage or not engage in it (*t*_(429)_ = 1.13, *p* = 0.13).

#### 3.2.5 Preliminary results into neurobiological basis of age-related increase in deliberation

Our behavioral analyses of the parallel experiments confirmed that the Dlx-CB1-KO mouse line reproduced the most central dimension of our aged mouse groups; deficit in EdM performance accompanied by increased rather than decreased deliberation (supplementary figure S.1B-D; Kruskal-Wallis or ANOVA comparisons with Wilcoxon or Tukey post-hoc tests, Dlx-CB1-KO vs WT; decision latency, *H* = 166, *p* < 0.0001; VTE, *F*_(1, 4768)_ = 134, *p* < 0.0001; run-time, *H* = 183, *p* < 0.0001). Furthermore was the observation that, even more pronounced than in aged mice, Dlx-CB1-KO animals displayed no significant overall effect of trial difficulty in either VTE or run-time (figures 1C-1D; ANOVA or Friedman repeated measures tests; VTE, *F*_(4, 56)_ = 0.9, *p* = 0.4; run-time, *F*_(4, 54)_ = 0.31, *p* = 0.9). In decision latency also, though Dlx- CB1-KO mice did display a significant overall effect of trial difficulty (*F*_(4, 54)_ = 7, *p* = 0.0001), when level 0 trials were excluded from analysis on the basis of our ‘rehearsal’ hypothesis (see above and ‘Rehearsal’ section of Discussion) this significant effect entirely disappeared (*F*_(3, 40)_ = 0.7, *p* = 0.6). This is a particularly interesting result since, based on the Dlx-CB1-KO literature, we should not expect these animals to be impaired in rehearsal-like behaviors, nor would such behaviors per se be impacted by a specific active forgetting deficit.

An interesting contrast between the aged populations and these young Dlx-CB1-KO animals is also worth highlighting. The Dlx-CB1-KO mice did display significantly increased VTE compared to WT animals, unlike aged compared to young animals. This suggests the possibility that the lack of increase we observed in aged mice may have resulted from an age-related decrease in physical vigor manifest in comparatively reduced full-body VTE behavior (see Materials & Methods). Regarding the WT animals from these parallel experiments, as with their EdM performance profile seen above, we observed that they robustly and in some cases even more significantly replicated all three deliberation behavior profiles of our baseline ‘Young*ITI10s’ population.

### 3.3.Anticipatory deliberation / deliberation as function of trial outcome

When we further analyzed the three deliberative behaviors according to ultimate per trial choice outcome (correct/rewarded versus incorrect/unrewarded; figure 4), we observed marked behavioral traces of what we interpret to be neurocognitive anticipatory processes (Steiner & Redish, 2014; Sweis et al., 2018). Indeed, our prediction had been that the lower number of EdM errors observed in young mice would be accompanied by distinct deliberative behavioral traces of anticipated trial outcome, while the RIF deficit putatively underpinning the age-related EdM impairment would conversely lead to surplus competing noise, anticipatory uncertainty, and thereby less or no such clear outcome-based discrimination observable in deliberative behaviors. On the basis of our tentative hypothesis that the Dlx-CB1-KO mice represent a model for cognitive active forgetting ‘knock-out’, we furthermore predicted that, compared to aged animals, they would display at least equal absence of deliberative discrimination as a function of trial outcome.

#### 3.3.1 Anticipatory discrimination decreases as a function of age

According to our graphic representations of the data, trial outcome appeared to have least impact on decision latency (figure 4A). Confirming this, paired Friedman repeated measures tests within each ‘Group*ITI’, revealed a slightly significant effect of outcome only in the ‘Aged*ITI30s’ population (*F*_(1, 5)_ = 9, *p* = 0.03). Moreover, whether this particular significant difference is the result of random variance within the very high variability aged population data (clearly visible in figure 4A, right) or the result of a genuine emergent cognitive impact of longer ITI values on aged mice is a question which will require further replication to satisfactorily answer. Furthermore, regarding the ‘Young*ITI10s’ population, although analysis did not show it to be statistically significant here, the small apparent increase in decision latency visible on low difficulty incorrect trials (figure 4A, left, red dashed vs red solid line) was replicated, in this case highly significantly, in the WT population from the Dlx experiments (supplementary figure S.1B, red dashed vs red solid line; *F*_(1, 17)_ = 34, *p* < 0.0001). This replication result strongly indicates that the statistically non-significant emergent outcome-based difference observed in the ‘Young*ITI10s’ population does have some underlying cognitive significance, despite manifesting to a more subtle extent here than in the WT animals.

With respect to VTE, where we had observed no significant difference in overall quantity of deliberation between young and aged animals (figures 2B and 4B), a two-way ANOVA nevertheless revealed not only a significant main effect of ‘Outcome’ (*F*_(1, 4476)_ = 16, *p* < 0.0001; further confirmed by a paired repeated measures ANOVA within ‘Outcome’, *F*_(1, 47)_ = 10, *p* = 0.002) but also a significant ‘Group*Outcome’ interaction effect as well (*F*_(1, 4476)_ = 9, *p* = 0.002). However, when we conducted paired repeated measures ANOVAs within each ‘Group*ITI’ population, we found that the effect of ‘Outcome’ reached statistical significance only in the ‘Young*ITI10s’ population (*F*_(1, 15)_ = 8, *p* = 0.01, effect size *η²* = 0.1). A clear yet visibly smaller and not quite statistically significant trend towards a similar impact of ‘Outcome’ was observed in the ‘Young*ITI30s’ population (*F*_(1, 15)_ = 3.5, *p* = 0.08, effect size *η²* = 0.04). By contrast, no such trend or significant effect was observed in either Aged*ITI population.

To explore the ‘Group*Outcome’ interaction in VTE in more depth, we calculated normalized delta values (correct- incorrect/correct+incorrect) for the aged and young animals (pooling across ITI values to focus on Aged vs Young differences). We then analyzed these values using a one-way ANOVA comparison with Tukey post-hoc test which confirmed that, as suggested by the graphic representations, VTE in young mice was significantly more likely to be correct-outcome oriented than in aged mice, indicating a task performance significant level of underlying accurate anticipatory representation (supplementary figure S.2A; *F*_(1, 567)_ = 12, *p* = 0.0005). Indeed, we were further able to reinforce this interpretation by conducting one-tailed T-tests between the delta values for each group and the 0 level around which VTE behavior would be expected to fall if it had no accurate reward-location anticipatory dimension. These again confirmed that only young mice were significantly more likely to engage in correct-outcome oriented (or rectifying) than to engage in incorrect-outcome oriented (or error-inducing) VTE deliberation behavior (T-tests with Welch correction; ‘Young’ vs 0, *t*_(317)_ = 7, *p* < 0.0001; ‘Aged’ vs 0, *t*_(250)_ = 1.4, *p* = 0.08). The significant difference between the delta values and 0 was also replicated in the WT animals from the Dlx-CB1-KO experiments (*t*_(250)_ = 12, *p* < 0.0001). We must nevertheless draw attention to the peak and accompanying jump in variability of error-inducing VTE in the ‘Aged*ITI30s’ population on level 2 difficulty trials (figure 4B right, indigo curves). This mean value peak is the result of two extreme outlier values and presents at least the possibility that, in replication, a certain level of anticipatory VTE outcome discrimination could be observed in aged mice at longer ITIs, as also suggested by apparent VTE outcome discrimination on trials of levels 0-1 only in the same group.

Finally, with regards to run-time (figure 4C), repeated measures Friedman comparisons with one-tailed Wilcoxon post-hoc tests confirmed the trends apparent in our graphic representations as well as our prediction that, in young but not in aged mice, there would be a significant difference in run- time as a function of whether ultimate trial outcome was unrewarded (incorrect) versus rewarded (correct) (*F*_(1, 29)_ = 14, *p* < 0.0001; *F*_(1, 13)_ = 1, *p* = 0.55, respectively). To explore this behavior in more depth, we calculated the ratio of incorrect- to correct-outcome run-times in both aged and young animals (independently of ITI). A one-way ANOVA comparison with Tukey post-hoc test revealed a significant difference between the respective ratios, demonstrating a significantly higher incorrect-outcome run-time bias in young compared to aged mice (supplementary figure S.2B; *F*_(1, 1762)_ = 5, *p* = 0.02). Subsequent T-tests between these ratio values and the incorrect/correct balanced value of 1 did however reveal that aged animals also displayed a slight but significant overall incorrect-outcome run-time bias, albeit with a noticeably smaller effect size than in young mice (Aged, *t*_(781)_ = 4.3, *p* < 0.0001, Cohen effect size, *d* = 0.15; Young, *t*_(981)_ = 9, *p* < 0.0001, *d* = 0.29).

#### 3.3.2 Putative interaction between ITI and VTE anticipatory deliberation

Returning to the difference observed in the effect sizes of ‘Outcome’ on VTE behavior in young animals as a function of ITI, we further separately analyzed the normalized delta values of the ‘Young*ITI10s’ versus ‘Young*ITI30s’ populations (data not shown) to see if the latter was indeed significantly lower than the former. However, a one-tailed T-test revealed only an almost significant trend in this direction (*t*_(300)_ = 1.5, *p* = 0.08). Nevertheless, combined with the difference in ‘Outcome’ effect sizes between the ‘Young*ITI10s’ and ‘Young*ITI30s’ populations noted above, the trend confirms that this is another interesting question to be revisited in future replications.

#### 3.3.3 Interaction of trial difficulty and anticipatory deliberation

Of great cognitive interest was the observation that the trial difficulty-related decrease in deliberative behaviors also took on a clear outcome-dependent profile, most notably in VTE and run-time (figure 4B-4C). Firstly, in young mice, we measured a highly significant impact of ‘Difficulty’ on correct-oriented VTE (repeated measures ANOVA, *F*_(4, 124)_ = 5, *p* = 0.005) but no such impact on incorrect-oriented VTE, which maintained consistently low values across all difficulty levels (*F*_(4, 124)_ = 1.3, *p* = 0.3). Conversely, in run-time, we observed a highly significant impact of ‘Difficulty’ on incorrect-oriented run-time lengths (repeated measures Friedman test, *F*_(4, 144)_ = 8.5, *p* < 0.0001) but no impact on correct-oriented run-time lengths, which again remained consistently low across all difficulty levels (*F*_(4, 144)_ = 2, *p* = 0.1). Indeed, the phenomenon observed may be best described as post-incorrect choice run-times decreasing as a function of increasing trial difficulty, down to an apparent minimal action execution time/maximal level of response vigor, comparable to that seen on post-correct choice trials at all levels of difficulty, including the easiest. The longest run-times in young mice thus occurred precisely on the level 0 trials where the very memory episode upon which anticipatory regret must be based is at its most recent and so, we would expect, most vivid also. The suggestion then is that, in young mice: anticipatory regret (of the kind that may also sometimes cause a subject, via VTE, to rectify its decision) is representational; elicits hesitative deliberative behaviors additively with respect to a baseline observed to be unaffected by varying levels of EdM challenge, and; progressively loses behavior-impacting power as a function of repeated RIF.

#### 3.3.4 EdM performance and anticipatory function in aged mice

In order to further investigate the relationship between anticipatory behaviors and EdM performance, we conducted a further subpopulation analysis on our data. First, we divided both our young and aged mice populations (independently of ITI), according to their overall mean EdM performance, into low performers (mean < 60%) and high performers (mean >= 60%). When we graphically represented the data from the small number of aged mice thereby classed as high performers (*n*=4; 2 animals from each ITI population), their outcome-based anticipatory deliberation profiles tended towards those observed in young mice (supplementary figure S.3A-3C). However, with the group of interest representing such a small sample size, statistical power for meaningful analysis was lacking, and so these graphically visible trends can serve only as invitations to further replication.

#### 3.3.5 Preliminary results into neurobiological basis of age-related loss of anticipatory accuracy

As has already been detailed, WT animals from the Dlx-CB1- KO experiments almost perfectly replicated each of the characteristic outcome-dependent deliberative behavior profiles observed in the ‘Young*ITI10s’ population. We have also already seen that the Dlx-CB1-KO animals reproduced in extremis the aged mice profile of EdM performance deficit accompanied by overall increased levels of deliberative behavior, and that these behaviors (in the case of VTE, run- time, and decision latency at levels 1-4) were similarly even less impacted by trial difficulty than in aged mice. Such similarity was again seen when we analyzed the impact of ‘Outcome’ on the deliberative behaviors observed in the Dlx- CB1-KO experiments: Dlx-CB1-KO animals displayed an exaggerated version of the profiles seen in aged mice, which is to say no effect of ‘Outcome’ or almost total absence of accurate anticipatory cognition in all three deliberation behaviors (repeated measures Friedman or ANOVA; decision latency, *F*_(1, 12)_ = 0.6, *p* = 0.4; VTE, *F*_(1, 14)_ = 0.1, *p* = 0.7; run- time, *F*_(1, 12)_ = 0.6, *p* = 0.4).

**Table 1.** Detailed summary of how experimental conditions relatively impact EdM performance and deliberation.

|  | YOUNG |  |  | AGED |  |
| --- | --- | --- | --- | --- | --- |
| ITI | 10s | 30s | AGED:<br>YOUNG | 10s | 30s |
| Errors | baseline | ↑<br>(*p* = 0.001) | ↑<br>(*p* = 0.001) | baseline | ↔<br>(*p* = 0.9) |
| Decision Latency | baseline | ↓<br>(*p* = 0.04) | ↑<br>(*p* < 0.0001) | baseline | ↔<br>(*p* = 0.36) |
| Decrease effect of 'Difficulty' | Yes*<br>(*p* < 0.0001) | Yes<br>(*p* = 0.003) | n/a | Yes<br>(*p* = 0.0006) | No<br>(*p* = 0.6) |
| Anticipatory Trace | Yes*<br>(*p* = 0.14) | No<br>(*p* = 0.3) | n/a | No<br>(*p* = 0.99) | Yes<br>(*p* = 0.03) |
| VTE | baseline | ↓<br>(*p* = 0.06) | ↔<br>(*p* = 0.9) | baseline | ↑<br>(*p* = 0.15)<sup>†</sup> |
| Decrease effect of 'Difficulty' | Yes*<br>(*p* = 0.15) | Yes<br>(*p* = 0.009) | n/a | No<br>(*p* = 0.2) | No<br>(*p* = 0.5) |
| Anticipatory Trace | Yes*<br>(*p* = 0.01) | Yes<br>(*p* = 0.08) | n/a | No<br>(*p* = 0.8) | No<br>(*p* = 0.9) |
| Run-time | baseline | ↔<br>(*p* = 0.7) | ↑<br>(*p* < 0.0001) | baseline | ↔<br>(*p* = 0.7) |
| Decrease effect of 'Difficulty' | Yes*<sup>‡</sup><br>(*p* = 0.0003) | Yes<br>(*p* = 0.05) | n/a | Yes<br>(*p* = 0.006) | No<br>(*p* = 0.45) |
| Anticipatory Trace | Yes*<br>(*p* = 0.05) | Yes<br>(*p* = 0.01) | n/a | No<br>(*p* = 0.18) | No<br>(*p* = 0.9) |
| * Effects replicated with statistical significance in WT animals from the supplementary Dlx-CB1-KO experiment (supplementary figure 1). |  |  |  |  |  |
| † Presence of significant opposing ITI impacts as function of age corroborated via bootstrap resampling approach. |  |  |  |  |  |
| ‡ Highly significant impact of 'Difficulty' observable only in run-times following incorrect choices (figure 4C). |  |  |  |  |  |

## 4. Discussion

Our investigation into the cognitive processes underpinning everyday-like memory (EdM) function has revealed a previously undescribed relationship whereby deliberation over representational mnemonic content observably decreases as a function of increasing EdM trial difficulty, the nature of which is precisely to entail increasing occasions for the amnesic effects of active- and retrieval induced-forgetting (RIF). We have further observed that trial outcome-dependent effects on the extent of deliberation are driven additively through anticipatory regret *increasing* deliberative behaviors via hesitation rather than through anticipatory satisfaction decreasing it via amplified vigor. With respect to the impact of ageing, we have discovered that the previously described EdM deficit of aged animals is accompanied by significantly *more* deliberation, largely independent of trial difficulty, and yet of significantly *lower* anticipatory accuracy. However, this is also just what we should expect to observe if the representational ‘stuff’ of mnemonic deliberation does undergo significantly less amnesic effect in aged compared to young animals, since accurate anticipation in the EdM task is reliant on successfully identifying and inhibiting (actively forgetting) the earlier of two directly competing memory episodes. Hence, our observations add significant weight to the hypothesis that EdM deficit in aged animals results primarily from an age-related impairment in specifically active forgetting processes. Aged mice deliberate more because they forget less, but because they forget less, they cannot accurately anticipate or select. As a general principle— though active forgetting does ultimately decrease performance and levels of deliberation as EdM challenge become more difficult—when active forgetting is impaired, EdM performance is severely negatively impacted across all trial difficulty levels, including the easiest.

Beyond the inherent interest for various domains of investigation that these novel behavioral results hold in and of themselves, cross comparison of the distinct deliberative profiles we were able to observe across five experimental conditions also opens the way to a broad program of future research. In this perspective, the following detailed discussion section will include a global interpretation of EdM function, comprising a range of hypotheses and predictions regarding its underlying cognitive and neural processes, plus proposals of literature-relevant experimental means by which these hypotheses can be tested in future research.

### 4.1 Active Forgetting & Everyday-like Memory

Our starting point interpretation is that EdM errors increase as a function of trial difficulty primarily as a result of the repeated instances of inhibitory active- and retrieval-induced forgetting (Bekinschtein et al., 2018; Tanaka et al., 2019; Costanzi et al., 2021; M. C. Anderson & Hulbert, 2021; M. C. Anderson et al., 1994) entailed by the continuous mnemonic retrieval and organization the EdM protocol was designed to elicit (Al Abed et al., 2016). Since it has been demonstrated that this kind of retrieval-induced forgetting (RIF from now on) comprises a genuine and lasting amnesic effect (Hulbert et al., 2016), we have hypothesized that a mnemonic representation of *A*_*n-1*_ retrieved at the onset of a high difficulty trial *A*_*n*_ will be significantly more weakened (via repeated RIF) than a similar representation retrieved at the onset of a low difficulty trial. Importantly, this entails that mnemonic weakening is not a result of merely being retained in episodic memory (EM) for longer durations of time. Rather, it occurs because, during that time, the memory episode has been repeatedly elicited, then actively inhibited without being acted on, all as part of the ongoing processes of EdM selection and organization. Cognitively speaking, a memory episode corresponding to the action taken on a trial *A*_*n-1*_ necessarily constitutes interfering content (i.e. noise) that must be inhibited if it intrudes into the representational deliberation space during the mnemonic deliberative processes elicited by any interposed trials on pairs *B* and/or *C* prior to onset of trial *A*_*n*_. This representational deliberation space can be thought of either in terms of a global workspace/GW (Dehaene et al., 2003; Dehaene & Changeux, 2011; Mashour et al., 2020) or as working memory/WM (Lustig & Jantz, 2015).

A critical factor in EdM performance is the persistence of and resultant interference from memory episodes corresponding to choices made on *n-2* trials. The literature refers to the particular challenge these memories represent as ‘proactive interference’ (Marighetto et al., 2011; Al Abed et al., 2016; Dulas & Duarte, 2016). Unlike memory episodes related to pairs *B* or *C*, an *A*_*n-2*_ memory episode will be spatially relevant to pair *A* though episodically irrelevant to the correct action to be executed on a given trial *A*_*n*_ (since this action should always be the opposite to the choice made on trial *A*_*n-1*_). This entails that proactive interference requires deliberation of a more strictly episodic rather than merely spatial nature, i.e. more properly related to spatio*temporal* organization allowing for disambiguation between the respective memories of what was done in a given place ‘last time/episode’ and what was done in the same place ‘the time/episode before that’. The primary role of active forgetting of spatial context-irrelevant content might even be to create the cognitive conditions allowing attention to be focused on the subtler challenge of deliberation over these spatial context-relevant episodes. However, the two phases of deliberation are not unrelated since, the weaker signals become as a result of repeated RIF during pair *B* and/or *C* trials, the less identifiably strong the required *A*_*n-1*_ signal will be compared to the competing *A*_*n-2*_ signal and, as a result, the closer performance on *A*_*n*_ will approach random chance level (i.e. 50% errors). Ultimately we advance that repetition of RIF-induced mnemonic representational weakening gives rise to a diminished signal [*A*_*n-1*_ memory episode] to noise [*A*_*n-2*_ + *B*_*n-1*_ + *B*_*n-2*_ + *C*_*n-1*_ + *C*_*n-2*_ memory episodes] ratio, thereby increasing the likelihood of a decision error on *A*_*n*_. This opens the way to an information theory-based cognitive hypothesis for why EdM errors increase as a function of increased numbers of interposed trials (i.e. trial difficulty level). Every time a signal is trial-inappropriately represented, it is susceptible to being identified as noise, consequently actively inhibited, and thereby weakened by the amnesic effect of RIF (see supplementary table S.1 for a tentative schematic representation of these hypothesized processes). However, in the absence of RIF to adjust the ‘volume’ of the various episodes, disambiguation between signal and noise will be critically impaired.

Hence, with respect to the impact of ageing on EdM performance, since the animal and human literature points towards both active forgetting functionality and RIF diminishing with age (Al Abed et al., 2016; Marighetto et al., 2011; Ossher et al., 2013; Lustig & Jantz, 2015), the model we have outlined predicts a dichotomous cognitive scenario whereby, on a given trial *A*_*n*_, an aged animal will experience: 1) *More* sustained interference from irrelevant memory episodes corresponding to previous trial actions, both on the other pairs (e.g. *B*_*n-1*_, *C*_*n-1*_, etc.) and on the same pair (e.g. *A*_*n-2*_), yet for the same reasons; 2) *Less* prior RIF amnesic effect on the *A*_*n-1*_ memory episode that is required for successful performance on *A*_*n*_. These two dimensions conspire to generate more cognitive ‘noise’. In short, we should predict aged mice to be impaired both in inhibiting spatial context- *irrelevant* memory episodes from pairs *B* and *C*, and in episodically disambiguating between the two spatially *relevant* memory episodes, *A*_*n-2*_ versus *A*_*n-1*_. We should further expect the *A*_*n-2*_ memory episode to continue competing ‘for attention’ in WM even during execution of the correct *A*_*n-1*_- based action choice (and vice versa with an *A*_*n-1*_ memory episode during execution of an incorrect *A*_*n-2*_-based choice). This double-edged situation of persistent competing interference noise could explain why, in contrast to young mice, the proportion of EdM errors in aged mice reached an only slightly better than random chance performance asymptote beginning at trials of difficulty level 1. Indeed, perhaps more demanding of explanation is how, given this significant active forgetting deficit, aged mice actually manage to commit fewer errors at level 0.

Although still susceptible to competing interference from spatially irrelevant memory episodes, the essential difference between levels 1-4 and level 0 EdM trials is that only the latter *can* be successfully performed without contribution from the CA1 region of the dorsal hippocampus, something our team has previously demonstrated using lesion studies (Marighetto et al., 2011). These past results, supported by our present observations, suggest a functional redundancy at play in level 0 trials only, whereby either hippocampal EM encoding and subsequent retrieval *or* cortical WM retention can be employed to resolve the mnemonic decision (for a computational implementation and demonstration of this EM vs WM redundancy, see Zilli & Hasselmo, 2008). By contrast, for performance at all trial difficulty levels above 0, on the hypothesis that WM will have been ‘cleaned’ of all content irrelevant to the spatial context of the interposed trial(s), recourse to EM is taken to be necessary. In other words, subsequent to an interposed trial, a given *A*_*n-1*_ memory episode will become accessible only via retrieval, and thus deliberation over it will in turn become susceptible to any cognitive deficits in retrieval-related functions, such as RIF (Bekinschtein et al., 2018), that a given subject (e.g. aged, lesioned, transgenic, etc.) may possess. This suggests that EM retrieval processes are in themselves sufficient to significantly increase the probability of irrelevant memory intrusion and competition (Wimber et al., 2015), whereas decision-making on level 0 trials may or may not have recourse to retrieval from EM (see ‘Replay’ section below). Hence, as long as aged mice are simply retaining something in WM, they do seem capable of keeping competing interference suppressed to a significant extent, something which has also been observed in human WM ageing studies (Staub et al., 2013). However, since an animal never knows what trial is coming next, this does not mean that level 0 trials are fully ‘protected’. During the intertrial interval (ITI), if an animal engages in cognitive activity such as replay, then the immediately previously executed action choice may be ‘pushed out’ of WM and encoded into EM, in which case, even on a subsequent level 0 trial, the appropriate memory will no longer be present in WM and will instead have to be retrieved from EM, thus incurring all related RIF deficit effects, etc., discussed above. Indeed, this may well explain the significant performance difference not only between young and aged mice on level 0 trials but also between young mice performing at ITI10s vs ITI30s. (Note also that the existing literature on aged phenotypes in simple reinforced T- or Y-maze spatial alternation protocols is inconclusive, some studies showing no significant impairment compared to young mice (Bisaz et al., 2013) some showing a highly significant deficit (Matzel et al., 2011), a discrepancy which may be due to differences in experimental design.)

In the preliminary supplementary study accompanying this paper, we nevertheless observed that Dlx-CB1-KO mice (animals lacking cannabinoid type-I receptors on all GABAergic neurons of the forebrain, Monory et al., 2006, investigated here as a putative model of active forgetting knock-out) revealed themselves to be incapable of inhibiting competing interference, giving rise to a generalized failure of EdM performance across all difficulty levels, including level

0. This is an even more important finding than it may at first appear, since it demonstrates that memories of previous spatially irrelevant actions do require active inhibition so that WM can successfully deliberate over context relevant memory episodes. The reason this is clear is that the lack of WM deficit previously noted in Dlx-CB1-KO mice (Albayram et al., 2016; a result we furthermore replicated in the radial maze using a single pair Y-maze configuration; data not shown) has been demonstrated only using simple T- or Y-maze configuration tasks, having no ‘other pairs’ and hence no necessary requirement for the inhibitory control or EM retrieval functions of WM. In other words, the experimental contexts used in previous studies were overly limited in terms of task environment-related interfering content, and so may simply have obfuscated a deficit in the nevertheless key active inhibition dimension of WM. By contrast, the EdM protocol continuously mobilizes this dimension, even on ‘Y-maze equivalent’ level 0 trials. These findings should further give us pause to consider what it means to investigate and conclude about WM function under experimental conditions which effectively exclude this active inhibition dimension.

### 4.2 Neurocogntive Processes of EdM Function

Evidence from the human literature has demonstrated that adaptive forgetting manages mnemonic interference via suppression of representational patterns at the level of the prefrontal neocortex (Wimber et al., 2015), making this region a putative locus for the conscious cognitive crossroads modeled as the GW or WM (but see Mashour et al., 2020 for a neuronally distributed conception of GW having no brain region ‘locus’ per se). In turn, this suppression from the prefrontal cortex has been shown to increase tonic GABAergic activity in the hippocampus, downregulating hippocampal activity related to the suppressed memory item (Malik et al., 2022; Schmitz et al., 2017). This is therefore one strong candidate neural process which may underlie the representational weakening putatively constitutive of RIF.

Perhaps the most common approach employed in the spatial EM literature to date is *in vivo* electrophysiological recording of place cells in the CA1 region of the hippocampus in freely moving, task-performing animals, though a calcium-imaging equivalent approach is now also possible (for an example of the latter, see Hazon et al., 2022). In a different radial maze protocol, (Xu et al., 2019) used an electrophysiological approach to demonstrate that during WM trials whose success relied on not repeating previous actions, CA1 cells in rats held in the central platform preferentially replayed previously executed context-related choices, i.e. those choices they must not repeat. Xu et al. advance that this replay of previous choices is a cognitive strategy that is specific to the WM task, since in reference memory tasks they instead observed ‘prospective’ activation representing animals’ intended future choice. Using a double Y-maze apparatus also in rats, (Ainge et al., 2007) observed more CA1 place cells activating when the animals were in the starting block compared to when they were in other locations, indicating a form of ‘mental time travel’ (Johnson et al., 2009; Redish, 2016) replay of all previously acquired trajectories related to the starting block context (4 in total in this case, as compared to 6 related to the central platform in our task). Finally, previous studies using either delayed non-match to sample (DNMS, another protocol where action must alternate from decision to decision) or EM tasks have observed that weakened CA1 activation was correlated with a higher probability of error (Hampson & Deadwyler, 2000; Ferbinteanu & Shapiro, 2003, respectively). In the EdM protocol then, to test the hypothesis that memory episodes – those related to the global radial maze environment but spatially or temporally irrelevant to a given trial – may intrude and competitively interfere during post-trial onset deliberation, CA1 activity could be recorded over repeated training in order to identify place cells specifically corresponding to all 6 radial maze arms used. Once identified, their activation during deliberation on trials to which they are irrelevant could then be measured. If our hypothesis regarding repeated active forgetting and consequent amnesic effect is accurate, we predict that trial relevant (i.e. *n-1*) place cell activation at a trial *n* will be weaker as a function of increasing trial difficulty. On a more physiologically grounded cognitive level, we predict this relationship will be tighter still if analyzed as a function of increasing numbers of effective prior intrusions and inhibitions, as identified from monitoring of actual brain activity during deliberation on interposed trials. Moreover, with such an approach, we also stand to learn which mnemonic contents are more likely to be replayed or retrieved outside of their context-relevant trials and thereby actively forgotten/subject to RIF; the most recent *n-1* memories or the earlier *n-2* ones? In this sense, Xu et al. 2019 were able to show that past WM task trajectories replay in reverse order, indicating that replay does display some manner of recency preference (note however that the WM task in question was not an active forgetting-dependent task). Hence, if spontaneous context-irrelevant replays and intrusions of the more recent *n-1* trial memory episodes are similarly more likely in the EdM protocol, this would entail that once a memory episode is shunted to *n-2* status by the formation of a corresponding *n-1* memory episode, *n-2* would be subject to little to no *additional* RIF. The representational strength of the *n-1* memory episode will then be weakened by RIF towards or even beyond the level of RIF-induced weakening previously reached by the *n-2* episode. To illustrate, if *n-2* was a difficulty level 1 trial and *n-1* a level 4, then the *n-2* representation will have been susceptible to RIF for 3 trial lengths *less* than the *n-1* representation. By this interpretation, when both *n-1* and *n-2* episodes are retrieved as spatial context-relevant episodes at onset of trial *n*, competition between their relative levels of RIF-induced weakening will be a critical determining factor in selecting which of them actually corresponds to the most recent decision action. Hence, in the example, the *n-2* level 1 trial memory, despite being *older*, would be *less* weakened by RIF than the more recent *n-1* level 3 memory. In future work, we will use a computational approach to test the predictions of such a model (laid out in preliminary schematic fashion in supplementary table S.1).

Similarly, what impact does the outcome of a trial have upon if and how often it is replayed? Such hypotheses could only be satisfactorily tested using the kind of high resolution method that either *in vivo* electrophysiology or calcium imaging recordings in the CA1 can offer. Thus, once again, Xu et al. 2019 observed that error outcomes did disrupt replay in rats performing in the radial maze. By contrast, in the present study, we were able to look only at the impact on post-trial onset deliberation (where we observed no significant impact of *n-1* outcomes on magnitude of deliberation on subsequent corresponding trials *n*; data not shown). With regards to aged and Dlx-CB1-KO mice, we would similarly predict that the active forgetting deficit will be visible in comparatively *less* weakening of all potentially interfering memory episodes on a given trial, including the proactively interfering memory episode *n-2*. In other words, we would predict a higher level of generalized, trial context-irrelevant CA1 activity.

#### 4.2.1 Endocannabinoid System and EdM Function

The close similarities observed between the aged and Dlx- CB1-KO mice phenotypes suggest the possibility that the active forgetting deficit in the former may be strongly linked to previously identified physiological age-related decrease in volume of CB1 receptors and subsequent impact on GABAergic processes (Albayram et al., 2011; Bilkei-Gorzo, 2012). Convergent evidence from other recent studies also points in the direction of a major eCS contribution to EM function. Notably, chronic exposure to a CB1 agonist (THC) restored memory reversal performance (being a type of cognitive flexibility putatively requiring a basic level of active forgetting) in aged mice in a Morris water maze task (Bilkei-Gorzo et al., 2017). Replication of such a THC protocol in aged mice performing the EdM protocol could directly test whether improved memory function in aged mice following chronic THC stimulation of the eCS is specifically the result of a rejuvenation of active forgetting capacities. It would also be of great interest to locally re-express CB1 receptors in GABAergic neurons of, respectively, the prefrontal cortex and hippocampus of Dlx-CB1-KO mice in order to evaluate the contribution of these region specific CB1 populations to active forgetting and EdM performance. If fruitful, this could be translated to a further step wherein CB1 receptors could be locally overexpressed in aged mice in order to test whether such an increase in CB1 volume is sufficient to recover some or all of the young animal EdM functionality.

### 4.3 Deliberation Behaviors

We turn now to discussion of the principal addition the present study brings to the literature, being the characterization of deliberation profiles arising from performance, under various experimental conditions, of an active forgetting-dependent EdM protocol. Taken together, the three deliberative behaviors we have classified (decision latency, vicarious trial-and-error/VTE, and run-time) occupy a temporal window that opens at the instant of trial onset and closes at the instant of arrival in the distal zone of the definitively chosen arm. Within that window, decision latency and VTE behaviors interact and overlap, in the sense that any VTE which occurs will happen within what is concurrently being quantified as decision latency, i.e. the time elapsed between trial onset and the instant of crossing the arm threshold on the definitive choice. This instant of definitive choice also delineates the end of decision latency from the beginning of run-time. However, with regards to VTE and run-time, although they too are similarly quantitatively separated by the same definitive choice instant, we interpret them as being qualitatively interrelated under the hypothesis that the cognitive processes which give rise to an animal withdrawing from an arm entered but not fully explored (i.e. VTE as operationally defined in this study) likely overlap with the cognitive processes which give rise to the kind of hesitancy phenotype we have seen manifest in slower post-choice run-times. Within our model, hesitancy is taken to be a reflection of indecision regarding whether an as of yet unconsummated choice should be concluded or abandoned. Because of this intricate overlapping of the three parameters, we have chosen to interpret and discuss them according to their various contributions to more chronologically constitutive cognitive phases of the global mnemonic decision-making process, namely: rehearsal, replay, conflict resolution, and anticipatory satisfaction or regret.

First, a brief introductory note regarding how our analyses began to move in this direction. In young mice performing at the longer ITI of 30s, we had observed that pre-definitive choice deliberation behaviors tended to be less pronounced than their corresponding values at ITI 10s, namely in decision latency (significantly lower) and VTE (almost significant trend). This was true even on level 0 trials and was accompanied, especially on trials of difficulty 0 and 1, by a significant increase in errors. Considering this observation in light of the literature led us to introduce two auxiliary hypotheses into our model: 1) That animals facing EdM challenge engage in ‘rehearsal’ (“the effortful maintenance of information in working memory,” Basile, 2018) of immediately completed trial action during ITI, and; 2) That while retained in the central platform during the ITI, and having no way of predicting which task context they will be presented with next, animals also spontaneously engage in putatively random cortico-hippocampal ‘replay’ of recent previous action choices from all task contexts (Ainge et al., 2007; Xu et al., 2019). Note that both of these cognitive phenomena, rehearsal and replay, must occur in large part during the ITI itself and therefore cannot be directly accounted for within our three behavioral deliberation parameters. However, we will see in the following two sections that they nevertheless cast a shadow of observable behavioral consequences.

#### 4.3.1 Rehearsal

‘Rehearsal’ has been well characterized in humans in behavior such as (overt or covert) verbal repetition of, for example, a phone number that must be remembered (Basile, 2018). Similar mnemonic rehearsal behavior has also previously been observed in rodents (Panlilio et al., 2011; Tanaka et al., 2019). Indeed, in their radial maze task, (Xu et al., 2019) also observed that the action choice of a trial just completed is ‘encoded’ by CA1 cells in rats upon returning and being confined to the central platform. Although we did not conduct electrophysiological recordings in the present study, we propose a mnemonic rehearsal-based interpretation of one of our behavioral observations. Hypothesizing that mice do engage in mnemonic rehearsal following each completed trial, we posited that on level 0 trials specifically, onset of a trial *n* plus concomitant re-opening of the same pair of doors as on the immediately preceding *n-1* may sometimes occur while this cognitive process is still ongoing. In such circumstances, the specific modification to the animal’s immediate environment (doors open) would afford the possibility (Gibson, 1979) of physically recruiting the newly available relevant element (i.e. the arm that was just visited on *n-1*) to the service of the ongoing cognitive process of rehearsal, i.e. via a partial, physical retracing of steps along the arm chosen on the immediately preceding trial. We propose that this kind of retracing action can be understood analogously to verbal repetition rehearsal in humans (Basile, 2018). Anecdotally speaking, we had previously noted that mouse deliberation in the EdM protocol did sometimes appear as much like an intentional retracing of steps (such as a human might engage in as a means of refreshing their spatiotemporal mnemonic bearings in order to retrieve a mislaid item) as it did a strictly corrective or choice-revising behavior. In terms of measurement, any such physical rehearsal behavior still ongoing at trial onset would be operationally captured under the decision latency and VTE measures quantified under the level 0 bin. Now, according to this interpretation, lengthening the ITI would increase the likelihood of rehearsal behavior being ‘complete’ (covertly, in this case) prior to the doors opening at trial onset, thereby entailing less cognitive cause for environmental recruitment/physical retracing to take place. Indeed, we suggest this could explain why a trend towards lower decision latency and VTE on level 0 trials was observed at ITI 30s compared to ITI 10s in young mice. Rehearsal behavior as just described could also explain some of the highly significantly greater rectifying than error-inducing VTE observed at level 0, since rehearsal behavior would lead to animals initially retracing precisely that action they had just taken on the previous *n-1* trial, being the action of which they must now, on trial *n*, choose the opposite. In other words, initial movement down the ‘incorrect’ arm would be ‘revised’ into definitive movement down the correct arm, giving rise, by our operational definition of this behavioral parameter, to a rectifying VTE event. Finally, again pointing in the direction of our interpretation, the effect size of the difference between rectifying and error-inducing VTE on level 0 trials was also greater at ITI 10s than at ITI 30s.

Interestingly, in aged mice, there was a slight tendency only at level 0 for decision latency to be higher at ITI 10s than at ITI 30s, indicating that aged mice too may engage in such rehearsal behavior during ITI. This would reflect recent demonstration of intact WM rehearsal capacity in aged human subjects (Hering et al., 2019). Finally, even Dlx-CB1-KO mice displayed significantly higher decision latency on level 0 trials only. On the basis of the literature demonstrating that this mouse line displays no deficit in retention or simple WM, we can posit that they too may engage in normal rehearsal behavior and only begin to encounter cognitive difficulties once they have to deliberate over retrieved mnemonic cognitive contents.

#### 4.3.2 Replay

To begin, some background on replay-relevant particularities of WM tasks. Spatial alternation WM tasks are based on one of the most robust spontaneous behaviors identified in rodents and other animals; contingent on recall of their previous choice, on a subsequent trial animals will reliably tend to choose the foregone option from the prior trial (Dember & Richman, 1989; Deacon & Rawlins, 2006). In certain protocols, including our EdM model, this alternation behavior is reward reinforced, rendering it even more robust. In order not to repeat a previous action, one optimal cognitive strategy would be to recall to memory precisely that action which is to be avoided. Indeed, it has recently been demonstrated that hippocampal replay in a WM task in the radial maze preferentially represents previous choices rather than the choice the animal intends to make next (Xu et al., 2019). By extension, under a WM rule, immediate post-*n-1* trial rehearsal, if it occurs, and later pre-*n* trial replay (during either ITI or trial *n* deliberation) will have the same representational episodic object. The first particularity the EdM protocol adds to this situation is that animals must spatially alternate in three related but mutually irrelevant spatial contexts concurrently, entailing a multiplicity of more or less conflicting mnemonic content being replayed and organized.

A second important particularity, in contrast to the WM task used by Xu et al. (2019), is that from the animal’s perspective the sequence of trial contexts in the EdM task is random, meaning the animal cannot reliably ‘plan’ its next trajectory during the ITI: it does not know on which alternation task it is going to be tested next. We therefore suggest that, while the animal is contained in the central platform, random previous trial episodes will spontaneously replay, in turn randomly conferring a kind of predictive priming advantage. In short, if an intra-ITI spontaneous *A*_*n-1*_ replay happens to temporally occur just prior to onset of a trial *A*_*n*_, this should confer a performance advantage on that trial. However, under a plausible hypothesis that, during ITI, the cognitive process of shifting awareness from replay of one previous trial action to another entails a certain level of organizational active forgetting, we would expect memory episodes retrieved for replay to be subject to levels of RIF similar to those resulting from competing memory episodes being inhibited during within-trial deliberation. Although no previous RIF studies have investigated this particular in-task question of ‘replay-induced forgetting’, one recent human study has shown that items replayed during sleep (and therefore similarly not *acted* upon) are nevertheless susceptible to RIF (Joensen et al., 2022). Hence, if our hypothesis is accurate, then lengthening the time an animal must wait enclosed in the central platform (ITI) until onset of a given trial, e.g. *A*_*n*_, would increase the likelihood for any given *n-1* memory episode from pairs *B, C*, or *A* to: 1) spontaneously replay; 2) be subsequently inhibited when one of the other *n-1* items is spontaneously replayed, and; 3) be thereby impacted by RIF-like amnesic effects, giving rise to reduced representational strength (plus concomitant reduced deliberation behavior) at onset of its subsequent relevant trial *n*.

In strict EdM performance terms, the result of all this would be an increase in errors across all difficulty levels, including level 0, in line with what we have observed in young mice performing at ITI 30s. This tentative interpretation also invites more general reflection on the underlying cognitive relationship between memory retention and active forgetting, since it intimately intertwines the latter with more classical concepts of ‘passive forgetting’ or ‘decay’ (i.e. simple passage of time posited as a putatively sufficient factor for deterioration of memory episodes; for further discussion of this point, see Hardt et al., 2013).

In contrast to young mice, aged mice displayed no significant increase in EdM errors at ITI 30s compared to at ITI 10s, including at level 0. Since, as we have just seen, errors did increase in young mice at ITI 30s, one consequence is that young and aged mice performing at ITI 30s had more comparable EdM scores, albeit accompanied by significantly differing deliberation profiles (see ‘Conflict Resolution’ section below). In light of the replay RIF hypothesis just outlined, the lack of error increase in aged mice at ITI 30s on level 0 trials is particularly striking. Can it too be explained on the basis of globally diminished RIF, manifest here in the context of mnemonic replay? If this be the case, the lack of inhibitory control over memories retrieved for replay could reflect human studies in which strength of cued purely mental recollection was self-reported as being more vivid, i.e. richer, in aged than in young subjects (St-Laurent et al., 2014), a finding which would also fit with the greater levels of overall deliberation seen in aged animals. Or do aged mice simply engage in less replay during ITI in the first place? If this be the case, putatively because aged mice may be too ‘distracted’ by encoding details of their immediate surroundings (i.e. the central platform) to engage in as much replay as the more goal-fixated young mice (a working hypothesis extrapolated from observations in human studies; Biss et al., 2013), then this potentially represents an important difference in EdM cognitive strategy between young and aged mice. Future studies employing *in vivo* electrophysiological or calcium imaging approaches may be able to demonstrate which of these two possibilities reflects actual intra-ITI neural activity.

#### 4.3.3 Conflict Resolution

Returning to the level of behavioral analysis, our strongest evidence for RIF consists in the observation that both of our pre-definitive choice deliberation parameters (decision latency and VTE) displayed *decreased* rather than increased values as a function of *increasing* trial difficulty. Intuitively, one may suppose that more difficult trials should occasion more uncertainty and therefore more, rather than less, deliberation. Indeed, had that been the case, it would simply have been understood as the logical flipside of the reduced deliberation previously shown to accompany proceduralization of action decisions (Pezzulo et al., 2016; for a review see Redish, 2016). Similarly, if one were to model organizational EdM memory according to a ‘stack’ structure (a common data structure in computer science, analogous to a stack of plates wherein adding or removing of items occurs via a last-in-first-out policy, i.e. always from the ‘top’ of the stack), then deeper objects, e.g. memory items which had been repeatedly ‘pushed’ under newer content, should also require *more* effort to subsequently access (J. R. Anderson & Lebiere, 1998; Newell, 1990; for a review and critique of stack models of goal-directed memory function, see Altmann & Trafton, 2002). However, in a paradigm just like that proposed by Redish (2016), where cognitive deliberation is conceived of precisely as deliberation over representational mnemonic and sensory items, then as the representational ‘stuff’ of a memory episode (whatever, for the neural code metaphysicists, that stuff may actually ‘be’) *loses* substance via an amnesic effect (e.g. via downregulation of related hippocampal activity, as mentioned above; Schmitz et al., 2017), so will there be *less* to deliberate with when that episode is retrieved into the representational deliberation space by its specific trial-appropriate context. Also important to note is that, in the course of the EdM protocol, repeated EM and WM challenges cycle continuously in such a way that, although mice do gradually improve their performances, no proceduralization stage is ever reached at which robust performance differences between low and high difficulty trials are no longer observable. Indeed, this represents a significant operational strongpoint of the EdM model, opening unprecedented access to complex yet robustly replicable mnemonic processes continuously active at multiple discrete levels of cognitive challenge. The robustness of this decreased deliberation as a function of increased active forgetting phenotype is attested to not only by the primary and supplementary experiments presented in the present study, but also in forthcoming work that further exploits the EdM model to study bias during revision of a previously acquired rule (Stevens et al., 2023b).

The decreasing deliberation phenotype also fits with the global interpretation we propose here, predicting that, on a given high difficulty EdM trial *n*, CA1 place cells participating in the representation of the action taken on a trial *n-1* will be *less* active than on a low difficulty trial. It also indicates that, in young mice, spatially irrelevant interfering content is easily identified and expediently actively inhibited. We assert this on the basis of the fact that, for any high difficulty trial, e.g. *A*_*n*_, although the corresponding *A*_*n-1*_ and *A*_*n-2*_ memory episodes may be significantly weakened due to repeated RIF, there will always be more recent and therefore more richly representational content from pairs *B* or *C* which would plausibly cause observably lengthy deliberation *if* they were not being rapidly inhibited on the basis of being identified as irrelevant to the current spatial context. The fact that we observe deliberation decreasing as a function of increasing trial difficulty thus suggests that they are being rapidly inhibited (in young mice at least). Hence, we once again suggest that the *primary* function of actively forgetting spatially *ir*relevant memory episodes is to create sufficiently noise free cognitive conditions allowing for the more subtle episodic disambiguation between equally spatially relevant *n-2* versus *n-1* memory episodes.

This leads us to what is perhaps the most important finding from our results with respect to the impact of ageing on EdM function. As stated above, in young mice we observed that lower difficulty trial performance was characterized by *more* deliberation but *fewer* errors. Likewise, in young mice, *decreased* overall deliberation at ITI 30s corresponded to *more* errors overall. However, in aged mice, we observed significantly *more* overall deliberation compared to young mice and yet significantly *more* rather than fewer errors. Why this sharp age-based contrast in the relationship between deliberation and performance? We interpret the aged mice phenotype of greater deliberation firstly as cognitive evidence that they are not impaired in the retrieval *per se* of representational mnemonic content into WM. Instead, we present it as still further evidence that the primary impact of ageing on EdM is specifically related to inhibitory active forgetting capacities, those which allow for directed suppression of competing, spatially and/or temporally irrelevant mnemonic content. These capacities, however, are precisely those required for making appropriate and accurate decisions via resolution of conflict between competing memories of previous choices, actions, goals, episodes (M. C. Anderson & Hulbert, 2021). Indeed, we further suggest that, on a certain proportion of trials, aged mice may fail to resolve this cognitive conflict at all and instead end up moving down one of the two open arms rather than the other, we might say, by default, i.e. despite *ongoing* unresolved conflict between competing mnemonic episodes. Supporting this interpretation, when we look at the post-choice run-time parameter, which we take to be a reflection of choice confidence, this too was significantly higher in aged mice across all difficulty levels and independently of outcome (see below), suggesting high levels of unresolved cognitive conflict persisting even post-choice, up until the end of the trial when the animal ultimately discovers whether its ‘default’ response was correct or incorrect (constituting, moreover, an environmentally imposed resolution of the cognitive conflict). The implication here for our tentative cognitive model of EdM function would be that, in aged mice, competing memory episodes are not always actively suppressed from neocortical WM, which would in turn entail that corresponding CA1 pattern activities are not downregulated, or at least not to the same extent.

Similarly contrasting with young mice, decision-making deliberation values in aged mice were also not impacted by ITI length. That aged mice displayed no significant differences in deliberation between ITI 10s and ITI 30s again fits with our interpretation of normal retrieval yet subsequent lack of RIF. Finally, regarding the discrepancy between significantly increased decision latency but not VTE in aged mice compared to young mice, our proposed interpretation of this observation is that, while the former do engage in more decision-making deliberation, it is possible that a lower proportion of this manifests in aged mice in the kind of large body movements our VTE parameter was designed to identify and quantify. In other words, a more fine-grained video-based behavioral analysis, precise down to the level of head movements, may reveal that aged mice do also engage in more VTE than young mice, albeit less vigorously. Supporting this possibility is the observation that young Dlx-CB1-KO mice did engage in significantly more VTE than wildtype animals. However, similarly to aged mice, their VTE behavior was independent of outcome (see below), leading us to suggest that they also often ‘chose’ an arm by default, due to a similar active forgetting deficit-based incapacity to resolve cognitive conflict between competing mnemonic episodes retrieved into WM. Indeed, recalling that the Dlx-CB1-KO mouse line has been reported to display normal spatial mnemonic retention and WM phenotypes (Han et al., 2012; Albayram et al., 2016), their EdM deficit with accompanying increased deliberation invites the hypothesis that this genotype is possessed of a previously undescribed active forgetting deficient phenotype, so pronounced it may even constitute an effective ‘knock-out’ of this cognitive process.

#### 4.3.4 Anticipatory Satisfaction or Regret

In young mice, all three deliberative behaviors reflected choice outcome, in what we interpret as an anticipatory manner (Steiner & Redish, 2014; Sweis et al., 2018). Decision latency tended to be higher prior to incorrect choices; the majority of VTE was rectifying rather than error-inducing, and; run-time was significantly higher following incorrect definitive choices. These phenotypes were particularly significant on (but not limited to) lower difficulty level trials. What we advance is that, in young mice, these behaviors reveal a contrast between anticipation of satisfaction and an anticipation of regret, the latter giving rise to indecision and response hesitancy (high rectifying VTE and slower run- times). Both these ‘flavors’ of anticipation can be thought of in the classical terms of a reward-prediction error (RPE) paradigm (Schultz, 2016). Interestingly, both error-inducing VTE and post-correct choice run-time were observed to be stably *low* across all difficulty levels, whereas *rectifying* VTE and post-*incorrect* choice run-time both tended to peak on level 0 trials then decrease as a function of increasing trial difficulty towards the stable baseline values of their respective opposites. By contrast, in the aged and especially Dlx-CB1- KO mice populations, in VTE and run-time, we observed more or less stably *high* values of outcome-independent hesitancy/indecision generalized across all trial difficulty levels. What this suggests is a striking impairment in accurate anticipatory function (putatively due to persistence of unresolved conflict between competing cognitive contents as described above), rather than a deficit in anticipatory function *per se*. Rather, these two mouse populations appear to persistently—beyond the definitive choice point, right up until the trial-terminating outcome reveal—alternate anticipation of *both* possible outcomes.

Bringing our representational interpretation to these observations, we suggest that in young mice choosing the incorrect arm on level 0 trials, the competing trial-appropriate *n-1* representation is sufficiently strong to provoke pronounced hesitancy (slower run-time); often even strong enough to trigger physical revision of choice (rectifying VTE). On high difficulty incorrect choice trials, by contrast, the equivalent competing trial-appropriate *n-1* representation is significantly weakened (due to repeated RIF) and correspondingly *in*capable of provoking marked levels of hesitancy. It is precisely this observable relationship between decision errors, longer run-time, and rectifying VTE which we interpret as reflecting hesitancy or anticipatory regret. Moreover, it is logical to posit that on trials where animals do physically revise their initial choice, the moment of retreat from the arm not fully explored will have been preceded by a similar, albeit potentially stronger, sense of anticipatory regret, leading first to hesitancy and deceleration, and then ultimately to choice abandonment. Future research with more fine-grained behavioral analysis capacity could test this prediction.

Looking to previous literature, we find evidence that such anticipatory cognitive behaviors may encompass a neurophysiologically grounded affective dimension, related (in part at least) to amygdalar function (McDonald et al., 2004; McDonald and Hong, 2004). Translating this affective dimension into everyday language, the suggestion is that animals experience an emotionally grounded ‘feeling’ or ‘sense’ about whether they are in the ‘right’ or ‘wrong’ place, i.e. about whether they have made the correct or incorrect decision with respect to reward location on a given trial. According to the general claims of paradigms such as Jaak Panksepp’s ‘affective neuroscience’ or Antonio Damasio’s ‘somatic marker hypothesis’ (Panksepp, 1998; Bechara & Damasio, 2005), such affective intuitions can have a significant impact on cognition. Thus, as far as dissecting the relative contributions of representational cortico-hippocampal versus ‘non-representational’ affective mnemonic content goes, the fact that decision latency, VTE, and EdM performance were impacted by ITI length, yet post-choice run-time was not, could be further evidence in agreement with the literature cited above (McDonald et al., 2004; McDonald and Hong, 2004) that run-time constitutes the parameter which, in young mice at least, is most sensitive to this ‘non-representational’ mnemonic affect. What we might suggest as a hypothesis for future testing is that any RIF amnesic effect occurring during ITI replay would have less of an affective impact (since it culminates in neither reward nor absence of reward to weight affective evaluation processes) than RIF occurring in the process of within-trial deliberation, which terminates in (positive or negative) reinforcement via reward consumption or reward absence. The broader question here regarding the nature of a potential relationship between RIF and affective reinforcement has not yet been answered in the literature, though it is of great potential interest for active forgetting research and could be investigated via a wide range of experimental approaches.

We do not exclude that a putative anticipatory affective dimension may also contribute to the pre-definitive choice deliberation phase. Evidence in this direction is observable in the trend for longer decision latencies prior to incorrect choices in young mice at low ITI values. One range of circuits through which such an affective dimension of reward location prediction could potentially contribute to spatiotemporal decision-making are the extensive projections from the basolateral (or pallial) amygdala (BLA) to the entire hippocampal complex (Freese & Amaral, 2009). There is also ample evidence that cross-talk between cortical-subcortical loops implicates the basal ganglia in the processing of non-motor signals, notably contextual, appetitive, and aversive ones (for review see Pessoa et al., 2022). Then there is the well-characterized bidirectional communication between the BLA and orbitofrontal cortex (OFC), centrally implicated in reward memory-guided decision making processes (for review see Wassum, 2022). Finally, that orbito-striatal connections play an important role in action choice has been demonstrated many times (Gremel & Costa, 2013; Gremel et al., 2016; Renteria et al., 2021), constituting robust evidence of a second neural avenue of communication through which affective dimensions could contribute to decision-making, including at the level of motor action selection. Indeed, these same avenues could also constitute the neural bridge between anticipatory ‘wrong place’ affect and the processes underpinning reduced response vigor (e.g. slower run-times), which the literature also strongly identifies with the basal ganglia (Carland et al., 2019; Dudman & Krakauer, 2016; Morrison et al., 2017). We suggest that, in the context of the EdM protocol, the affective dimension serves as a reciprocal contributor to overall representational deliberation, present to a greater or lesser extent depending on the phase of the decision making process, but being most evident in the final phases, those which are quantitatively subsumed in our study under VTE and run-time. Indeed, Damasio and others suggest the idea that different representations (be they mnemonic or sensory in origin) will elicit distinct feelings, constitutive of their context-dependent significance (or meaning) to the organism (Damasio, 2012; Pulvermüller, 2013).

That the maximal observed magnitude of hesitancy was similar in young (at level 0) and aged (at all levels) mice is commensurate with literature showing that the amygdala, compared to the hippocampus or cortex, is a brain structure that is relatively resistant to age-related deterioration in both rodents and humans (Von Bohlen und Halbach & Unsicker, 2002; Mather, 2016). This resistance can, however, be both a blessing and a curse, as previously demonstrated in the case of decision-making in humans (Denburg et al., 2006), because of prefrontal cortical inhibitory control over amygdalar signals (Rosenkranz & Grace, 2001) *decreasing* with age, analogous with the age-related reduction in cortico-hippocampal inhibitory control posited to underpin active forgetting deficit. Taken as a whole, what this suggests as a model of the impact of ageing on EdM performance is diminished cortico-hippocampal RIF in the pre-choice deliberation phase subsequently accompanied (not replaced) by diminished cortico-amygdalar affective inhibitory control during the post-choice anticipatory phase. Since we have already suggested that reduced active forgetting in aged mice would also increase proactive interference on a given trial *n* from spatial context relevant *n-2* trial memory episodes, this provides a further complementary putative explanation for why aged mice do not display an outcome-dependent run-time phenotype similar to that of young mice: rather than experiencing a strong feeling that the reward is on the other arm, they would experience strong ‘*mixed* feelings’ that the reward could be on either this *or* the other arm. Hence, they advance with equally pronounced ‘hesitation’ following both incorrect and correct choices. Here again, we observed that Dlx-CB1-KO mice displayed a very similar phenotype to aged mice: significantly higher overall VTE and run-time compared to WT mice, yet in a trial difficulty- and outcome-independent manner. In short, Dlx-CB1-KO animals also manifested what we suggest is the result of strong ‘mixed feelings’ that the reward could be on either this or the other arm. Since, as we have just said, not only hippocampal but also amygdalar signals are under inhibitory control from the prefrontal cortex, then if these mechanisms are impaired by deletion of CB1 receptors from GABAergic populations of the forebrain, this represents a promising candidate neural mechanism for the run-time phenotype seen in both Dlx-CB1-KO and aged mice. Again, this is not to exclude the possibility of persistent representational interference, especially if choice is made by default, i.e. without active forgetting of spatially irrelevant competing memory episodes. But, with respect to hesitation, reduced response vigor, anticipatory regret, etc., are all elements of cognition which may also receive a significant affective, putatively amygdalar, contribution.

Since a small subpopulation of aged mice achieved EdM performances comparable to those of young mice, the possibility that this tracked with persistence of accurate anticipatory reward location discrimination in these animals presented itself and seems, though the result is only preliminary, to have been upheld by the data. This tentative finding echoes intriguingly with a finding from human studies that only those aged individuals *not* displaying impaired decision-making capacities produced normal psychophysiological outcome-anticipatory skin conductance responses, in contrast to decision-making impaired aged individuals who did not (Denburg et al., 2006). Related to neural basis hypotheses formulated above, in future work it would be interesting to quantify CB1 levels in high versus low aged performers to test the prediction that these levels should be higher in high performers. (Note also that a similar subpopulation division could not be conducted with the Dlx- CB1-KO experiment data as only 1 out of 15 KO animals achieved a mean EdM performance above 60%, further demonstration of the extent and robustness of their active forgetting deficit.)

Reliable cognitive measures for post-choice confidence are relatively sparse in the rodent and especially mouse literature (Carandini and Churchland, 2013; Hanks and Summerfield, 2017; Kepecs et al., 2008), despite this being a fundamental component of deliberative decision making processes. In the radial maze, this component is gained via an inherent spatial fact of the apparatus itself, which enables analysis of cognitive processes in mice as their choice is physically unfolding in both space and time, thereby capturing both the “doing and undergoing of the consequences of doing” phases of behavior (Dewey, 1916). This is not possible where choice execution is more discrete (e.g. lever press, nose poke, etc.) and opens up exciting possibilities for deeper *in vivo* investigation into the neural bases of post-choice confidence/hesitancy, including but certainly not limited to the cortico-amygdalar hypotheses articulated above.

## 5. Concluding Remarks

Our hope for the results presented here is that they act as a draft blueprint for a testable theoretical unification of our neurocognitive understanding of active forgetting and RIF, deliberative spatiotemporal goal-directed decision-making (and related CA1 place cell activity), impact of ageing on these processes, and the precise role of the eCS in the various real world components of overall episodic memory function. The terrain of such a neurocognitive theoretical unification would likely spread beyond the domains here listed. For example, associative generalization (Samborska et al., 2022) and interference might be fruitfully considered as two manifestations of one and the same primitive cognitive process, in the sense that the phenotype we observed here in aged and Dlx-CB1-KO mice under the label of an ‘active forgetting deficit’ could as easily be described in terms of a form of ‘overgeneralization’ (Beck, 1979; Garner & Dux, 2022) or in terms of a previously identified age-related *deficit* in pattern separation yet *reinforcement* of pattern completion (Stark et al., 2010; Wilson et al., 2006). Indeed, the original incidental associative learning study that inspired us to test the Dlx-CB1-KO mice (Busquets-Garcia et al., 2018) revealed a phenotype in this mouse line which consists in failure to disambiguate an associative generalization into its distinct component parts. Relative to potential causes of the inhibitory control deficit, further investigation will be needed to disambiguate between a deficit in inhibitory mechanisms per se and neurocognitive conditions potentially impairing the ‘targeting’ of inhibitory control, such as excessive pattern completion.

With respect to animal cognitive capacities more generally, the present work also adds to an ever-growing literature (Eichenbaum, 2004, 2016; Johnson & Redish, 2007; Konkel & Cohen, 2009; McKenzie et al., 2014; Steiner & Redish, 2014) highlighting observable cognitive behaviors that are difficult to account for unless one assumes that the animal models under investigation do generate, maintain, and organize representational cognitive content in a manner analogous to humans (philosophical debates over what mental or cognitive representation actually ‘is’, even in humans, notwithstanding). For example, precisely because of the observed cognitive signs of persistent hesitancy both before and during exploration of arms visited during the EdM challenge, it would be difficult to support a potential objection that aged mice act in such irresolvable situations unreflectively, e.g. merely by ‘instinct’. It would nevertheless be fruitful to entertain, in the spirit of comparative pluralism, the implications of non representation-based interpretations of the behavioral data we have presented here, whether those be computational or cognitive in nature.

As a final consideration, while our EdM protocol has been designed in such a way as to generate specifically conflictual memory episodes, reflective of common everyday-like human situations, there is evidence to suggest that *congruent* distraction can, by contrast, be of more benefit to aged than to young subjects (Weeks & Hasher, 2014). This is an important point to consider for at least two reasons: 1) It mitigates hasty interpretation of age-related changes in cognitive function as being in and of themselves a necessarily net negative development for the organism (Lustig & Jantz, 2015); 2) It raises questions for further investigation in domains relating to the evolution of human environments and societies. For example, in the environment(s) in which human and proto- human cognition first evolved and emerged, which contexts were more frequent or advantageous; those giving rise to ultimately conflictual or ultimately congruent ‘distraction’? In other words, though age-related changes in cognition may appear particularly disadvantageous to everyday navigation of our modern industrial and technological environment, there is no *a priori* reason to presume this was always the case in environments almost unimaginably far removed from our own. In much the same way, no one would suggest mouse cognition evolved in order to optimize performance on a radial maze task. Hence, any complete understanding of everyday-like mnemonic cognition must inquire in this evolutionary direction also.

## Author Contributions

C.S., G.M. and A.M. designed research; C.S., A.S.A., A.S., E.D., C.L., M.B., and F.R. performed research; C.S. and A.M. analyzed data; C.S. and A.M. wrote the paper.

## Acknowledgements

We thank all of the personnel of the Animal Facility of the Neurocentre Magendie for mouse care. Countless members of the stackoverflow.com community were also of invaluable help during development of the Python-based analyses. This work was financially supported by: Université de Bordeaux (MESRI doctoral contract to C.S.), Institut National de la Santé et de la Recherche Médicale, Centre National de la Recherche Scientifique, Conseil Régional d’Aquitaine.

## Supplementary Figures

**Supplementary figure S1.**
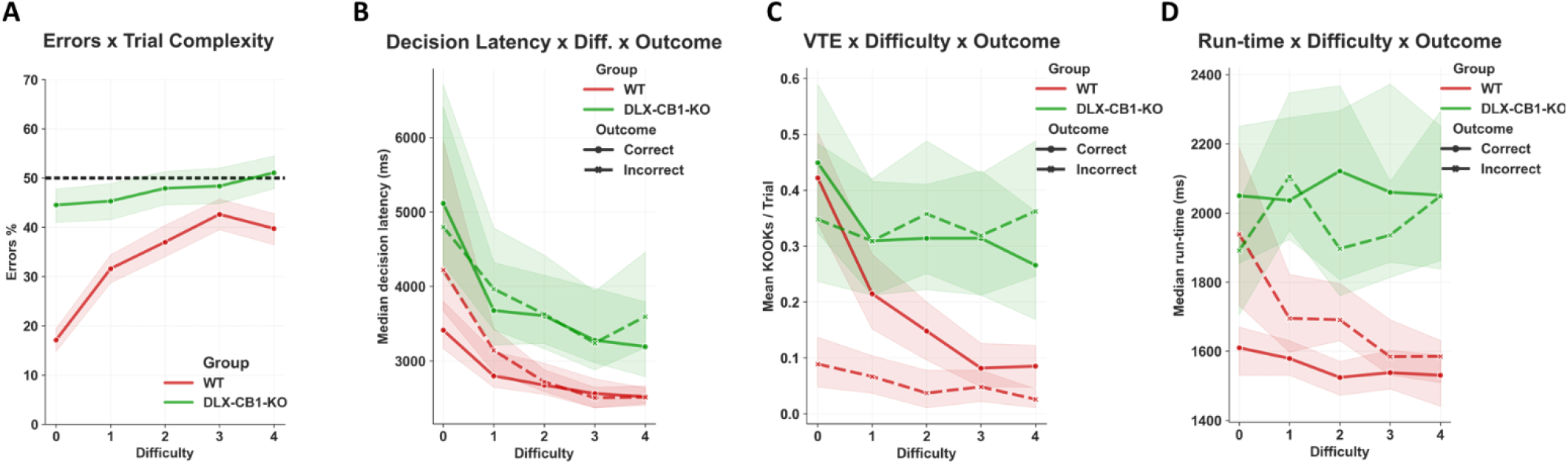
Dlx-CB1-KO as putative model of active forgetting knock-out. Experimental groups: Wildtype animals (WT, red curves; *n* = 20); Dlx-CB1-KO animals (green curves; *n* = 15). All experiments represented here were conducted at ITI values of between 3 and 10s. **A)** The profile of errors committed as a function of EdM trial difficulty by WT animals replicates that seen in the ‘Young*ITI10s’ population. Dlx-CB1-KO mice commit even more errors than aged mice, with only slightly better than random chance level performance even on level 0 trials, indicating a near total knock-out of EdM function. **B–D)** With respect to deliberative behaviors represented as a function of both EdM trial difficulty and trial outcome, WT animals again replicated the primary features seen in the deliberation profiles of the ‘Young*ITI10s’ group. By contrast, Dlx-CB1-KO mice deliberated significantly more, but with no observable impact of trial outcome, and in a less trial difficulty-dependent manner. Hence, in terms of both EdM performance and EdM-elicited deliberation, Dlx-CB1-KO mice display what can be described as an extreme version of the aged mouse phenotype, hinting at an almost complete cognitive knock-out of the cortico-hippocampal active forgetting capacity necessary for normal EdM function.

**Supplementary figure S2.**
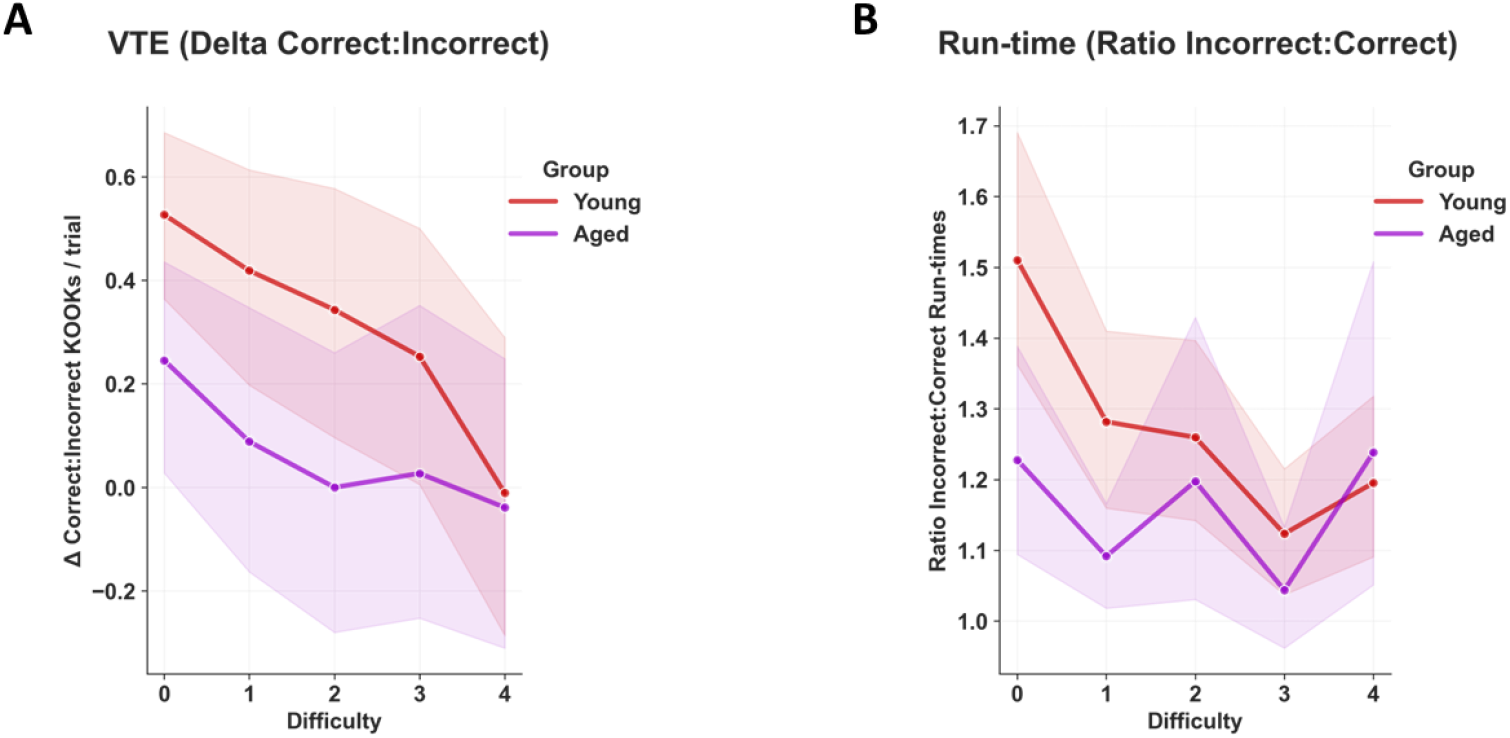
Deliberative behaviors normalized as function of trial outcome. Experimental groups (ITI populations pooled by age): Young (red curves; *n* = 32); Aged (indigo curves; *n* = 16). Results are averaged over multiple EdM training blocks, giving 12 to 48 trials per trial difficulty per animal. All error bands represent 95% confidence intervals; vertical spaces between bands provide visual indication of statistical significance. **A)** VTE delta values (correct-incorrect/correct+incorrect) for aged and young animals (pooled across ITI values) confirmed that overall VTE in young mice was significantly more likely to be correct-outcome oriented than in aged mice. **B)** Ratio run-time values (incorrect:correct) similarly confirmed that length of run-time in young mice was significantly more biased towards incorrect outcomes than in aged mice. See main text for details and compare with figure 4B-C.

**Supplementary figure S3.**
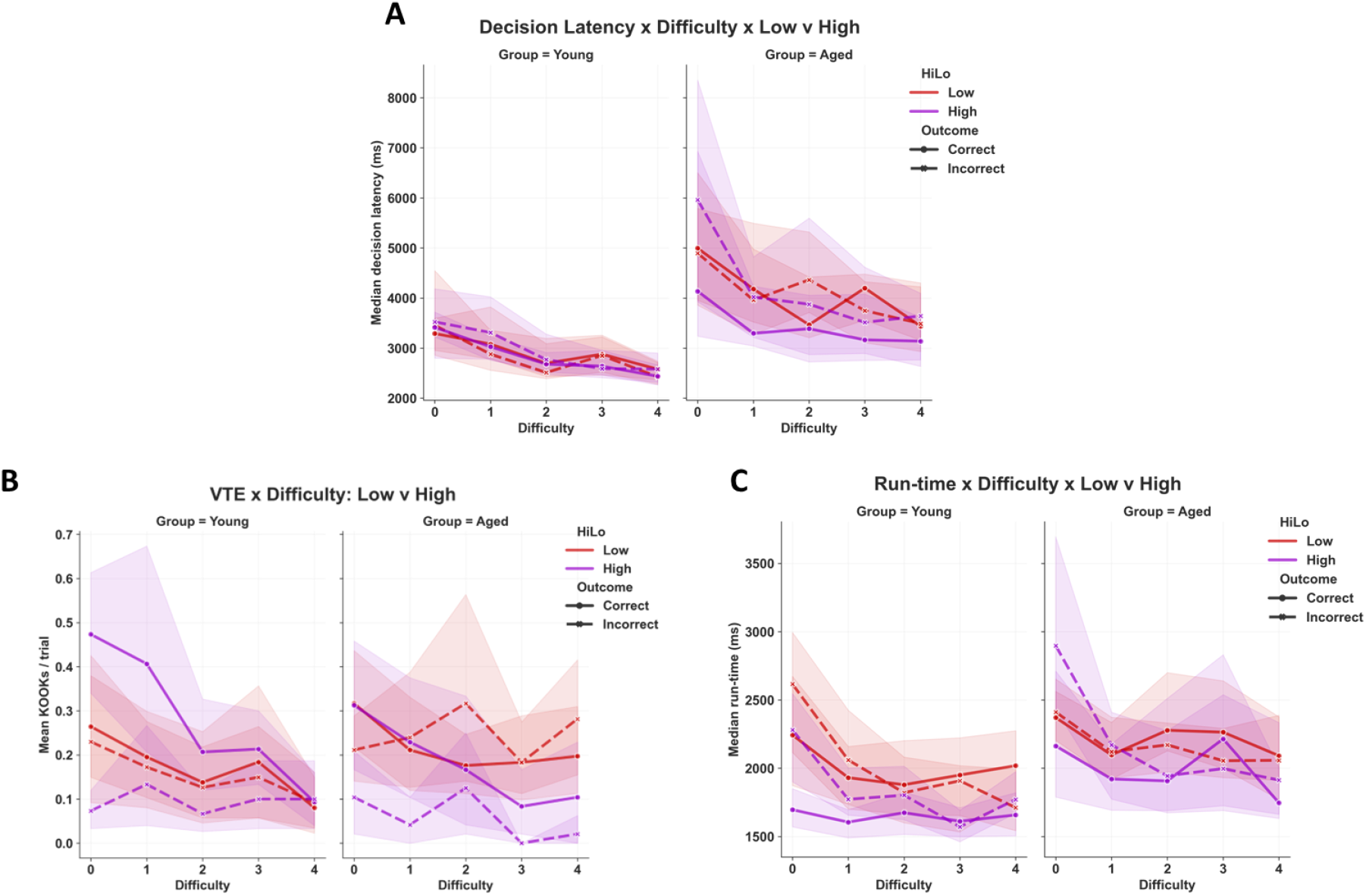
Deliberative behaviors displayed according to low versus high EdM performers. Groups (ITI populations pooled by age): Young*Low (left columns, red curves; *n* = 9); Young*High (left columns, indigo curves; *n* = 23); Aged*Low (right columns, red curves; *n* = 12); Aged*High (right columns, indigo curves; *n* = 4). Results are averaged over multiple EdM training blocks, giving 12 to 48 trials per trial difficulty per animal. All error bands represent 95% confidence intervals; vertical spaces between bands provide visual indication of statistical significance. Although the sample is small (*n* = 4), high EdM performance aged mice appear to display trial difficulty and outcome-dependent deliberation profiles similar to those seen in young mice.

**Supplementary Table S1.** *Schematic model of putative RIF impact on* n-2 *vs* n-1 *memories*. Here, we have presented mnemonic contents schematically to illustrate how more recent *n-1* memory episodes may be more susceptible to RIF than earlier *n-2* memory episodes. Notably, in the case of the trial designated **C’’**, it illustrates how an *earlier* memory, e.g. **C** (*n-2*), may nevertheless be *less weakened* than a more recent memory, e.g. **C’** (*n-1*), when both are elicited by the spatial context and need to be deliberated over. This highlights the kind of putative cognitive mechanisms by which an organism may mistake an earlier memory for a more recent one due to the earlier mnemonic episode being representationally richer, i.e. more vivid, than the more recent episode, due to the latter having been more repeatedly actively forgotten.

| Current Trial Context: | A | B | C | A' | C' | B' | A'' | A''' | B'' | C'' | ... |
| --- | --- | --- | --- | --- | --- | --- | --- | --- | --- | --- | --- |
| Memories elicited/retrieved (during trial): | n/a | n/a | n/a | A | C | B | A A' | A' A'' | B B' | C C' | ... |
| RIF factor '$n-2 : n-1$' representations: | n/a | n/a | n/a | n/a | n/a | n/a | 0 [-2:-2] | -2 [-2:0] | -1 [-3:-2] | 3 [-1:-4] | ... |
| Memories 'RIF'ed (during trial): | n/a | A | A B | C B | A' B | A' C' | B' C' | B' C' | A''' C' | A''' B'' | ... |
| New '$n-1$' memory (post trial):<br>(more susceptible to RIF) | | A | B | C | A' | C' | B' | A'' | A''' | B'' | C'' |
| Prior '$n-2$' memory (post trial):<br>(less susceptible to RIF) | n/a | n/a | n/a | A | C | B | A' | A'' | B' | C | ... |

Mnemonic episodes:
RIF factor color code: 0 1 2 3 4

